# Revisiting the trafficking of insulin and its receptor in rat liver: *in vivo* and cell-free studies

**DOI:** 10.1101/2023.09.07.498136

**Authors:** Khadija Tahiri, Françoise Fouque, Marie-Catherine Postel-Vinay, Bernard Desbuquois

## Abstract

Endosomes are the main locus where, upon endocytosis in liver cells, insulin and the activated insulin receptor accumulate and insulin is degraded. In this study, ligand and receptor pathways in rat liver have been revisited using analytical subcellular fractionation. Density gradient analysis of microsomal and light mitochondrial fractions confirmed that, upon *in vivo* uptake into liver, [^125^I]-insulin was rapidly translocated from plasma membranes to endosomes. Internalized [^125^I]-labeled proinsulin and biotinylated insulin, which are less degraded than insulin in endosomes, were in part transferred to lysosomes. Following injection of native insulin, receptor and ligand were also both translocated from the plasma membrane to endosomes, with a maximum at 15-30 min. Receptor translocation was reversed by 2-3 hours and neither endocytosis nor recycling were affected by *in vivo* cycloheximide treatment. Fractionation of endosomes of insulin-treated rats on density gradients showed that the insulin receptor was recovered at low density, whereas insulin progressively migrated from low to high densities towards the position of acid phosphatase. On SDS polyacrylamide gels, insulin receptor α and β subunits were identified in plasma membrane, endosomal and lysosomal fractions by affinity crosslinking and Western immunoblotting, respectively. Following insulin treatment, insulin receptor expression rapidly decreased in plasma membranes while increasing in endosomes. In a cell-free system, [^125^I]-insulin was in part transferred from endosomes to lysosomes, as was, with organelle content mixing, [^125^I]-biotinylated insulin. In summary: 1) the low lysosomal association of [^125^I]-insulin *in vivo* is linked to its endosomal degradation; 2) upon ligand-induced endocytosis, the insulin receptor progressively segregates from insulin and is recycled; 3) neither endocytosis no recycling of the receptor requires protein synthesis.

## Introduction

Liver expresses specific, high affinity insulin receptors which mediate insulin action as well as degradation. Using as ligand [^125^I]-insulin, liver insulin receptors were identified initially in plasma membrane (Freychet *et al*, 1971) and microsomal fractions (Cuatrecasas *et al*, 1971, 1972), and later in Golgi fractions (Bergeron *et al* 1973, Posner *et al*, 1978), nuclei and nuclear envelope (Goldfine & Smith 1976, Vigneri *et al*, 1978, Wong *et al* 1988), and isolated hepatocytes (Terris & Steiner, 1975, Olefsky *et al*, 1975). Shortly thereafter, morphological (Bergeron *et al*, 1977, 1979, Carpentier *et al*, 1979, Goldfine *et al*, s 1981) and biochemical (Desbuquois *et al*, 1979, 1982, Posner *et al*, 1980, Smith & Peters, 1982, Pease *et al*, 1985, 1987) studies showed that, upon *in vivo* receptor-mediated uptake into liver, [^125^I]-insulin undergoes endocytosis along with the insulin receptor. This process, mainly clathrin-dependent, involves the translocation of the insulin-receptor complex from the plasma membrane to endosomes (reviewed by Bergeron *et al*, 1985). Liver insulin receptors were also shown to undergo ligand-induced translocation from the cell surface to nuclei (Podlecki *et al*, 1987, Wong *et al*, 1988, Gletsu *et al* 1999, Nelson *et al*, 2011, Kesten *et al* 2018). Endocytosis of the insulin-receptor complex is implicated in insulin degradation, signaling and action (reviewed by Knutson, 1991a, Morcavallo *et al*, 2014, Bergeron *et al*, 2016, Petersen *et al*, 2018, Chen *et al*, 2019, Hall *et al*, 2020, Iraburu *et al*, 2021).

Instead of lysosomes, in which many proteins taken up into liver are degraded, endosomes are the main site at which, upon dissociation of the internalized insulin-receptor complex, insulin is degraded (reviewed by Authier *et al*, 1994, and Berg *et al*, 1995). Studies on cell-free systems have shown that [^125^I]-insulin degradation in liver endosomes occurs at low pH (4-5.5) and is functionally linked to ATP-dependent acidification by the vacuolar H(+)-ATPase proton pump (Pease *et al*, 1985, 1987, Smith *et al*, 1982, 1989, Doherty *et al*, 1990, Desbuquois *et al* 1990, 1992, Hamel *et al*, 1991, Bevan *et al*, 2000). Thus, degradation of [^125^I]-insulin in isolated endosomes is inhibited by ionophores (nigericin, monensin), weak bases (chloroquine) and ATPase inhibitors (N-ethylmaleimide). Owing to its ability to inhibit degradation, *in vivo* chloroquine treatment increases the association of [^125^l]-insulin with liver endosomes (Posner *et al*, 1982, Desbuquois *et al*, 1992) and extends the lifetime of the activated insulin receptor (Bevan *et al*, 1997). *In vitro* endosomal degradation of insulin is also inhibited by metal chelating agents (1,10-phenanthroline), thiol-blocking agents (p-chloro-mercuric benzoate) and the peptide antibiotic bacitracin (Peavy *et al*, 1985)

Degradation products of [^125^I]-insulin in intact liver endosomes (Hamel *et al*, 1988, Clot *et al*, 1990, Seabright *et al*, 1992) and of native insulin in endosomal extracts (Authier *et al*, 2001) have been isolated and characterized, and sites at which insulin A and B chains of are cleaved have been identified. Insulin degrading enzyme (IDE), a neutral thiol metallo-endopeptidase, has been implicated in insulin degradation in endosomes (Duckworth, 1988, 1998, Najjar, 2018, Leissring, 2021) and proposed to initiate this process when insulin is still bound to its receptor. However, liver-specific ablation of IDE in mice did not affect insulin clearance (Villa-Perez *et al*, 2018), and based on crosslinking, immune-depletion and immune-blotting procedures, IDE has been localized in cytosol and peroxisomes but not in endosomes (Authier *et al*, 1994, 1995, 1996). Instead, cathepsin D, a soluble pepstatin-sensitive aspartyl endopeptidase, has been identified as the major enzyme which initiates insulin degradation in liver endosomes (Authier *et al*, 2002)

As in cultured cells exposed to insulin *in vitro* (Kasuga *et al*, 1981), liver insulin receptor expression and subcellular distribution are regulated by plasma insulin *in vivo*. Upon acute hyperinsulinemia, ligand-induced endocytosis of the receptor in liver is rapidly reversed, total receptor number remaining unchanged (Desbuquois *et al*, 1982). In contrast, upon long lasting hyperinsulinemias, such as that observed in genetically obese Zucker rats, total cellular receptor number in liver is decreased while receptor number in endosomes is increased (Amessou *et al*, 2009). In Zucker rats, an increase in insulin receptor degradation accounts for the decrease in receptor expression (Amessou et al, 2009), as it does in isolated fibroblasts (Knutson, 1991b), and results in insulin resistance (Knutson *et al*, 1995). Ubiquitination of insulin receptor subunits by the E3 ubiquitin ligases c-Cbl (Ahmed *et al*, 2000), MG53 (Song *et al*, 2013) and March1 (Nagarajan*et al*, 2015), has been implicated in receptor degradation and inhibition of insulin signaling in liver and cultured cells. Unlike receptor ubiquitination by c-Cbl, which is induced by insulin, ubiquitination by March1 occurs in the basal state and results in a decrease in receptor expression at the cell surface. In the insulin-sensitive state, a decrease of March1 expression linked to inhibition of the transcription factor Foxo-1 results in an increased receptor expression. Some other negative regulators of insulin receptor expression are the adapter Grb10, which mediates ligand induced receptor ubiquitination and degradation (Ramos *et al*), and the β-secretase BACE-1, which cleaves the receptor ectodomain in the Golgi (Meakin *et al*, 2018)

As expected from the functional relationships between endocytosis and signaling (Sorkin & von Zastrow, 2009, von Zastrow & Sorkin, 2022), ligand-induced activation of the insulin receptor kinase and of downstream signaling pathways occurs both at the plasma membrane and in endosomes (reviewed by Baass *et al*, 1995, Bevan *et al*, 1996, and Bergeron *et al*, 1988, 1995, 2016). In both compartments, the activated receptor undergoes autophosphorylation at tyrosine residues (Khan *et al*, 1986, 1989) followed by dephosphorylation by phosphotyrosine phosphatases (Burgess *et al*, 1992, Faure *et al*, 1992, Drake *et al*, 1996, Li *et al*, 2006). Upon activation, the insulin receptor kinase recruits and phosphorylates a number of intracellular substrates that initiate discrete signaling pathways (reviewed by Siddle, 2011, 2012, Boucher *et al*, 2014, Saltiel, 2021). Major substrates are the IRS and Shc proteins, which initiate the activation of the PI3K/Akt and Ras/Raf/Mek/Erk signaling pathways, respectively; the former pathway is preferentially activated at the plasma membrane, and the latter in endosomes. IRS, p85 PI3K and Akt signaling proteins recruited by the activated insulin receptor undergo redistribution between these cell compartments (Balbis *et al*, 2000, Drake *et al*, 2000). A selective activation of endosomal insulin receptor kinase mediates the *in vivo* hypoglycemic effect of bisperoxovanadium, a phosphotyrosine phosphatase inhibitor (Posner *et al*, 1994, Bevan *et al*, 1995). Several proteins recruited by the insulin receptor, such as PC-1/ENPP1 (Goldfine *et al*, 2008, Youngren, 2007), SOCS proteins (Ueki et al, 2004) and Grb7/10/14 adaptors (Holt and Siddle, 2005, Desbuquois *et al*, 2013) inhibit receptor autophosphorylation or receptor tyrosine kinase, whereas others, such as SH2B1/B2 adaptors are activators. The cell adhesion molecule CEACAM1, which upon association with the insulin receptor is phosphorylated by the receptor kinase, stimulates the endocytosis of the insulin-receptor complex (Najjar *et al*, 2002, 2018). In doing so, CEACAM1 regulates insulin sensitivity by promoting hepatic insulin clearance (Poy *et al*, 2002). Insulin receptors translocated to liver nuclei along with insulin signaling proteins associate with the transcriptional machinery at gene promoters and regulate the expression of insulin-inducible genes (Nelson *et al*, 2011, Sarfstein & Werner, 2013, Hancock *et al*, 2019).

In most previous studies, trafficking of insulin and its receptor in rat liver *in vivo* has been characterized using isolated plasma membrane and Golgi/endosome fractions, which account together for 15-20% of total receptor content. In the present study, these trafficking events have been revisited using analytical subcellular fractionation, with focus on distribution of [^125^I]-insulin, native insulin and insulin receptor, physical characterization of the internalized receptor, and cell-free endosome to lysosome transfer of insulin and receptor.

## Results and discussion

### *In vivo* receptor-mediated uptake of [^125^I]-insulin into rat liver

Based on the rate of plasma disappearance and volume of distribution of combined high and low affinity [^125^I]-insulins, a receptor compartment with many properties of the receptor characterized *in vitro* has been identified *in vivo* in rabbits (Zeleznick & Roth, 1978). In subsequent studies, *in vivo* identification of insulin receptors in rat liver has been achieved by measurements of [^125^I]-insulin uptake into the intact organ (Sodoyez *et al*, 1980) and isolated subcellular fractions (Izzo *et al*, 1979, 1983, Desbuquois *et al*, 1979, Posner *et al* 1980, Gagliardino *et al*, 1980, Papachristodoulou *et al*, 1981, Pease *et al*, 1985, 1987). Insulin uptake into intact liver has also been assessed by electron microscope autoradiography (Bergeron *et al*, 1977, 1979, Carpentier *et al*, 1979) and scintigraphic scanning (Sodoyez *et al*, 1983, 1985, Sodoyez-Goffaux *et al*, 1987, 1988) following injection of [^125^I]- and [^123^I]-labeled hormone, respectively. As expected from a specific and saturable receptor-mediated process, uptake of [^125^I]-insulin was inhibited by excess co-injected native insulin but unaffected by unrelated hormones such as glucagon or growth hormone.

In agreement with a previous study (Posner *et al*, 1980), the ability of co-injected native insulin and insulin derivatives to inhibit uptake of [^125^I]-insulin was dose-dependent and in correlation with their *in vitro* binding affinities for the insulin receptor. The dose of native insulin required for half-maximal inhibition of [^125^I]-insulin uptake by a total liver particulate fraction was about 0.6 nmol, and des-alanine, des-asparagine insulin, proinsulin and des-octapeptide insulin were, respectively, 20%, 5% and 0.4% as potent as native insulin (supplementary Fig. 1).

Using an assay in which the *in vivo* uptake of injected [^125^I]-insulin into rat tissues was compared to that of [^125^I]-albumin, Whitcomb *et al* (1985) have shown that [^125^I]-insulin specific uptake was highest in liver. A similar observation has been made by Zeleznick and Roth (1978), who compared the *in vivo* uptake of high affinity chicken and low affinity guinea pig [^125^I]-insulins into rabbit tissues. In agreement with these studies, the *in vivo* uptake of [^125^I]-insulin into total particulate and microsomal fractions of rat tissues decreased in the order liver > adrenals > small intestine > lung > spleen > heart > adipose tissue > skeletal muscle (supplementary Fig. 2). As shown previously, kidney uptake of [^125^I]-insulin was as high as liver uptake but was not saturable.

### Subcellular distribution of [^125^I]-insulin taken up by rat liver *in vivo*

Preparative subcellular fractionation procedures have shown that, upon *in vivo* uptake into rat liver, [^125^I]-insulin is concentrated first at the plasma membrane and then in low density microsomal elements (Desbuquois *et al*, 1979, Posner *et al*, 1980). First proposed to originate from the Golgi apparatus, the latter were shown by the diaminobenzidine density shift procedure to be separable from the Golgi marker galactosyltransferase and hence identified as endosomes (Kay *et al*, 1984). On analytical subcellular fractionation, [^125^I]-insulin taken up into liver was time-dependently concentrated in several components of different density: a single high-density component in the nuclear (N) fraction, and two components (of low and high density) in mitochondrial-lysosomal (ML) and microsomal (P) fractions (Bergeron *et al*, 1986). Based on the distribution of marker enzymes, the low density component in ML and P fractions was assigned to endosomes, and the high density component in N and P fractions to plasma membranes.

Using analytical subcellular fractionation, we also identified a single component of high density in the N fraction and two incompletely resolved components in the ML fraction (results not shown), but only a single component in the P fraction. On sucrose density gradients, this component was time-dependently shifted from the positions of *in vitro* insulin binding activity and plasma membrane markers (peak density, about 1.16 g/ml) to those of galactosyltansferase and the mannose 6-phosphate/IGF-II receptor (peak density, 1.11-1.12 g/ml) (Fig 1). In agreement with previous studies (Pease *et al*, 1984, 1987), regardless of post-injection time digitonin treatment shifted the distribution of [^125^I]-insulin towards higher densities, at the position of plasma membrane markers, while shifting to a lesser extent the distribution of galactosyltransferase (Fig. 2, left panels).

**Figure 1.**
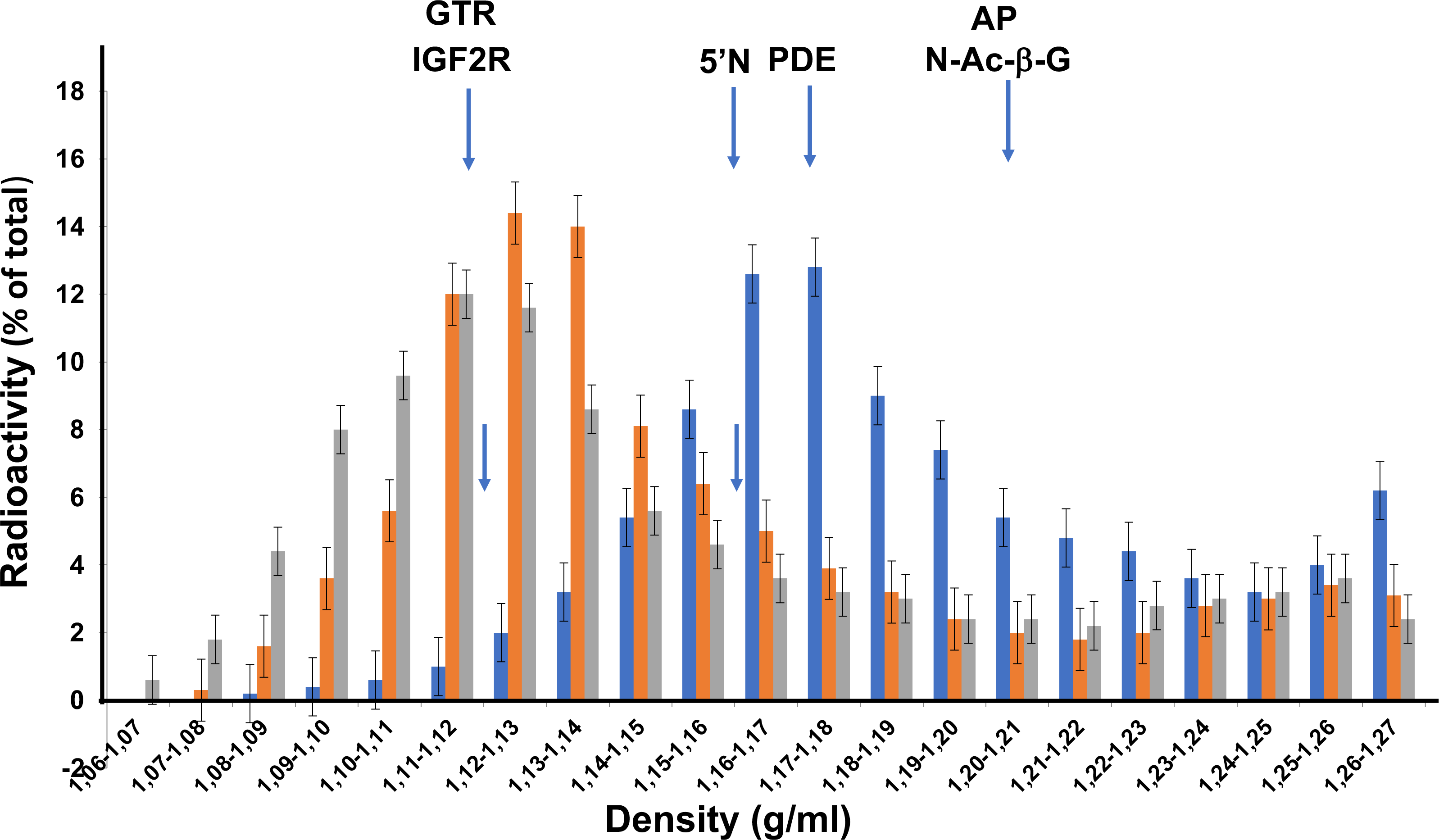
Density distribution of [^125^I]-insulin taken up into the liver microsomal fraction *in vivo*. Microsomal fractions were isolated at the indicated time after injection of [^125^I]-insulin and subjected to centrifugation on sucrose density gradient. The distribution of [^125^I]-insulin at 20 sec (blue bars), 90 sec (red bars) and 5 min (grey bars) after injection has been plotted against normalized density intervals of 0.01 g/ml. The results shown are the mean of 5 determinations. Arrows indicate the peaks of galactosyltransferase (GTR), IGF2/mannose 6-phosphate receptor (IGF2/M6P), 5’nucleotidase (5’N) and alkaline phosphodiesterase (PDE)

**Figure 2.**
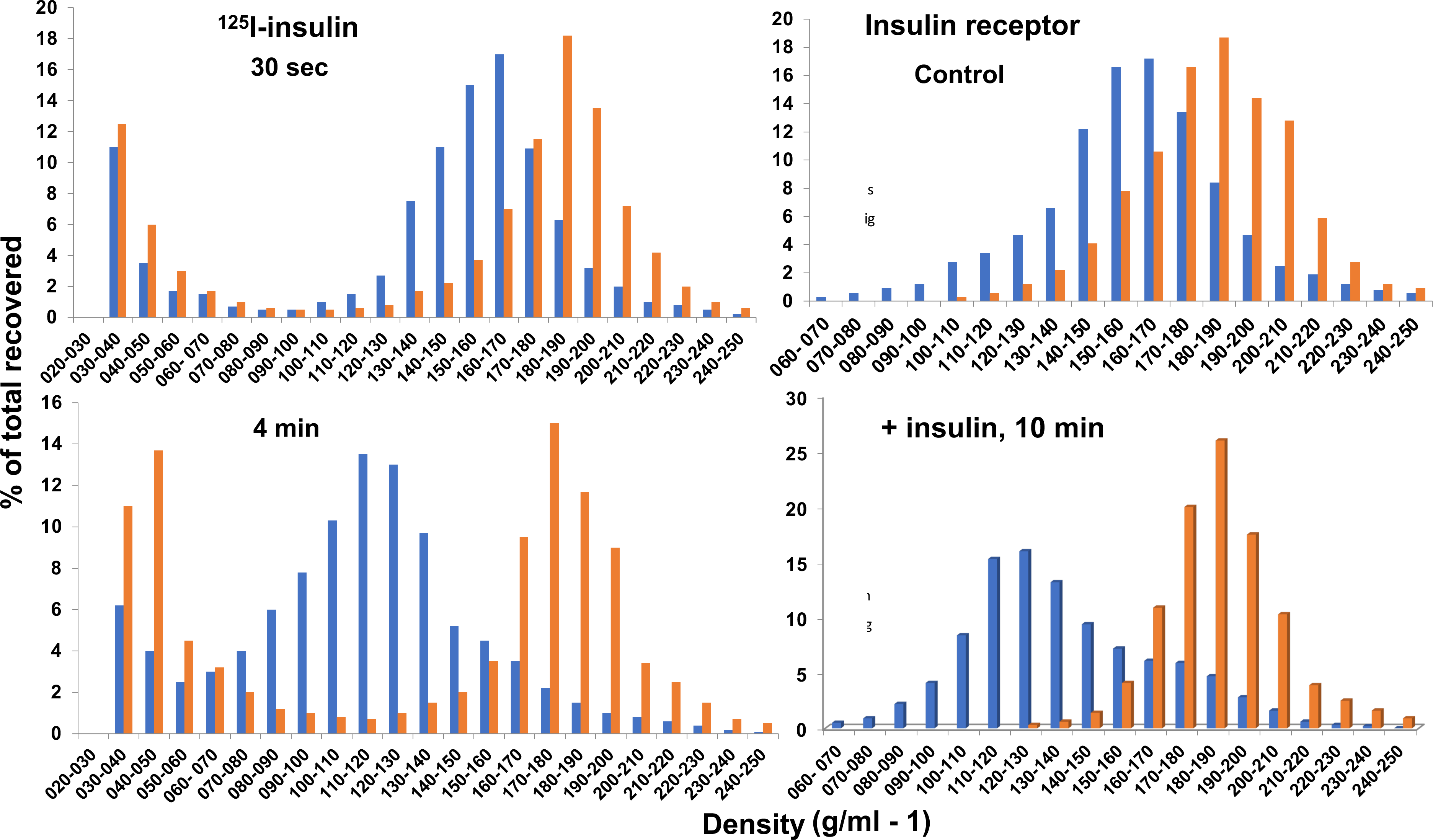
Effect of digitonin treatment of the microsomal fraction on the density distribution of injected [^125^I]-insulin and insulin receptor. Liver microsomal fractions were prepared at the indicated time after injection of [^125^I]-insulin (left panels) or native insulin (right panels); control rats (right panels) were left untreated. Fractions were treated by digitonin (red bars) or left untreated (blue bars), following which the distribution of [^125^I]-insulin (left panels) and of *in vitro* insulin binding activity (right panels) on sucrose density gradients was examined. Representative experiments (of three) are shown.

Although migrating in part at the position of lysosomal markers on density gradients (Bergeron *et al*, 1986), [^125^I]-insulin taken up into liver was recovered to a limited extent in purified lysosomal fractions, even after treatment by chloroquine (Posner *et al*, 1982). Furthermore, insulin-containing structures present therein, which contained lipoprotein-like particles, were considered as non-lysosomal (Posner *et al*, 1982). We have previously suggested that the low association of internalized [^125^I]-insulin with lysosomal fractions results from its rapid degradation in endosomes, resulting in the release of degradation products out of the endosomal lumen (Desbuquois *et al*, 1990, Chauvet *et al*, 1998). Accordingly, the distribution of [^125^I]-insulin associated with the L fraction on Nycodenz density gradients has been examined and compared to that of [^125^I]-proinsulin, which upon *in vivo* uptake into liver is degraded more slowly than insulin (Desbuquois *et al*, 2003), and of [^125^I]-biotinylated insulin, the α amino group of which is blocked. As shown on Fig. 3, the migration of the two latter polypeptides towards high densities, at the position of lysosomal markers, was markedly increased at 8 min and 16 min post injection relative to that of [^125^I]-insulin. These observations suggest that the low association of [^125^I]-insulin with the high density region of the gradient results mainly from its rapid endosomal degradation.

**Figure 3.**
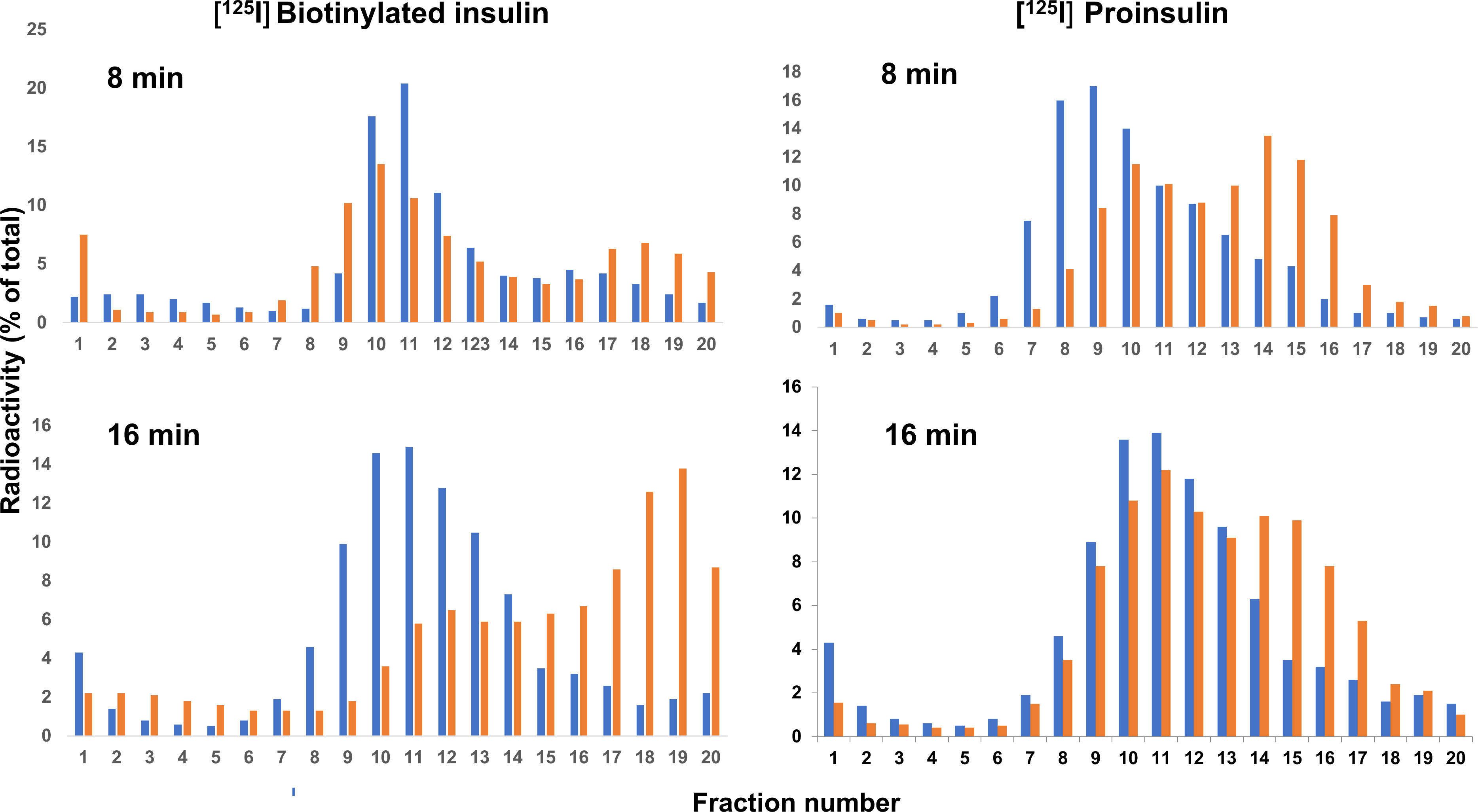
Comparative density distributions of [^125^I]-labeled insulin, biotinylated insulin and proinsulin taken up into light mitochondrial (L) fractions *in vivo.* L fractions prepared at the indicated time after injection of [^125^I]-labeled insulin, biotinylated insulin and proinsulin were sub-fractionated by centrifugation on linear Nycodenz density gradients (density range, 1.05 to 1.15 g/ml). Twenty subfractions were collected and assayed for radioactivity (blue bars, [^125^I]-insulin; red bars, [^125^I]-biotinylated insulin and [^125^I]-proinsulin). The results shown are representative of two to three experiments. N-acetyl−β-D-glucoinidase activity was recovered in fractions 12-20.

### *In vivo* insulin-induced changes in the subcellular distribution of the insulin receptor and of acid-extractable insulin in the microsomal fraction

We have previously shown that following injection of a high dose of native insulin to rats, *in vitro* insulin binding activity rapidly increases along with insulin content in Golgi/endosome fractions, while simultaneously decreasing in the plasma membrane fraction (Desbuquois *et al*, 1982). In later studies, these time-dependent changes of insulin binding activity and insulin content in Golgi/endosome fractions of insulin-injected rats have been confirmed and shown to be increased by chloroquine treatment (Khan *et al*, 1985, Desbuquois *et al*, 1992). In the present study, the insulin-induced changes in receptor and ligand distribution have been reexamined using analytical subcellular fractionation

The distribution of *in vitro* insulin binding activity and immunoreactive insulin in primary subcellular fractions was first examined (Fig. 4). In control rats, insulin binding activity was recovered mainly in the nuclear (N) and microsomal (P) fractions (about 30% and 65% of total activity, respectively). Following injection of a saturating dose of insulin, a time-dependent decrease in insulin binding activity in the N fraction and a reciprocal increase of activity in the P fraction were observed. These changes were detectable at 2 min, achieved maximum at 30 min and were later slowly reversed. Injection of insulin also led to a rapid increase of immunoreactive acid-extractable hormone in N and P fractions (about 3 and 60 ng insulin per g liver, respectively), followed by a slow, time-dependent decrease (half-life, about 30 min).

**Figure 4.**
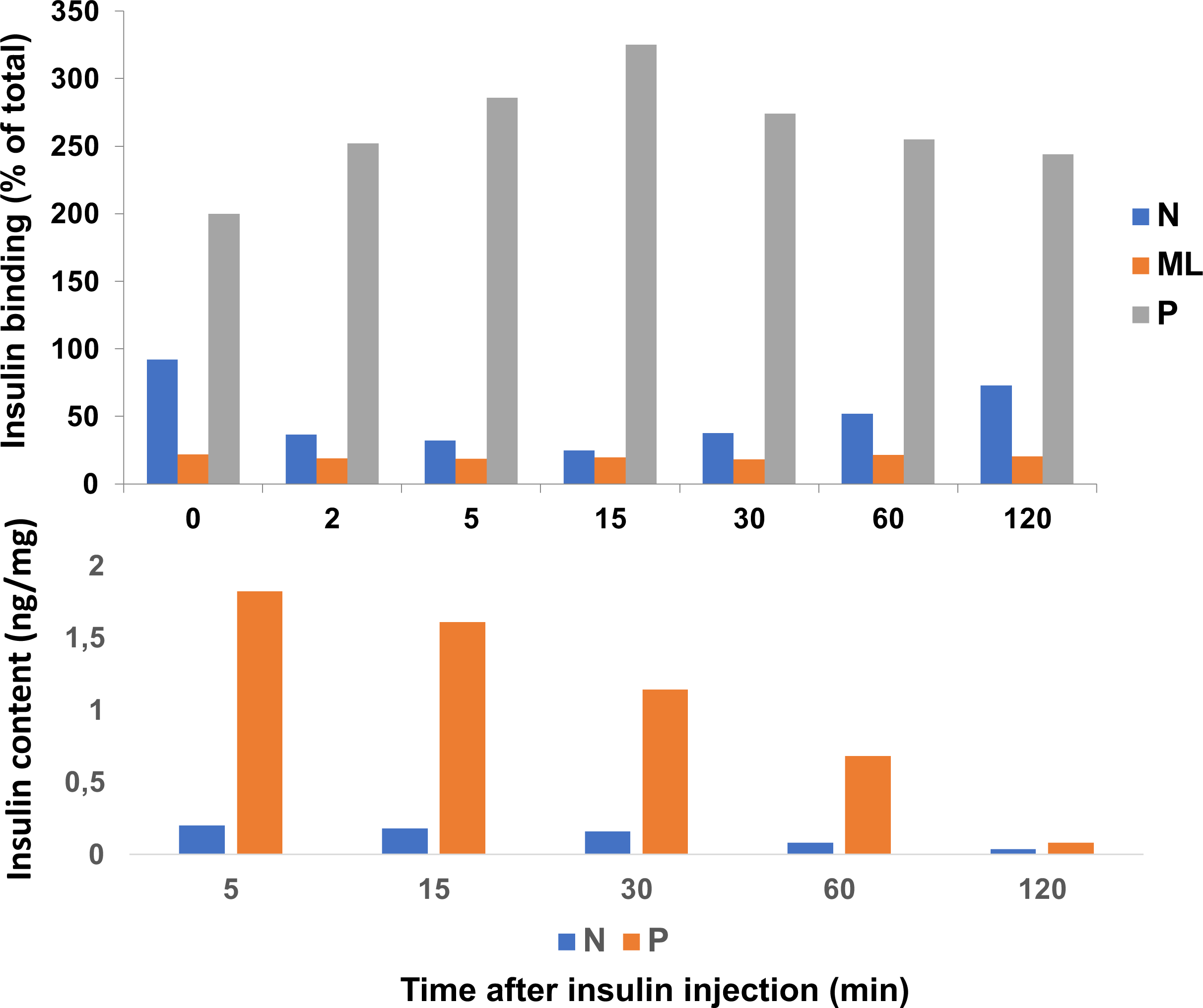
*In vitro* insulin binding activity and insulin content of primary liver subcellular fractions isolated after insulin injection. Rats were sacrificed at the indicated time after injection of insulin (50 μg) or left untreated. Liver nuclear (N), mitochondrial-lysosomal (ML) and microsomal (P) fractions were prepared and assayed for *in vitro* insulin binding activity and extractable immunoreactive insulin content. The results shown are the mean ± SEM of three determinations.

The distribution of *in vitro* insulin binding activity and extractable insulin in the microsomal fraction on sucrose density gradients was then examined (Fig. 5 and Table 1). In control rats, the distribution of insulin binding activity largely overlapped that of the plasma membrane markers 5’nucleotidase and alkaline phosphodiesterase (median density, 1.155 g/ml). Injection of insulin led to a time-dependent shift of insulin binding activity towards lower densities, which achieved a maximum at 15-30 min and was nearly reversed at 2 h. At 15 min, the density distribution of activity was similar to those of galactosyltransferase and the IGF2/mannose 6 phosphate receptor (median density, 1.115 g/ml). As shown above for [^125^I]-insulin distribution, digitonin treatment of the microsomal fraction shifted the distribution of insulin binding activity on sucrose gradients towards higher densities (median density, 1.179 g/ml) (Fig 2, right panels).

**Figure 5.**
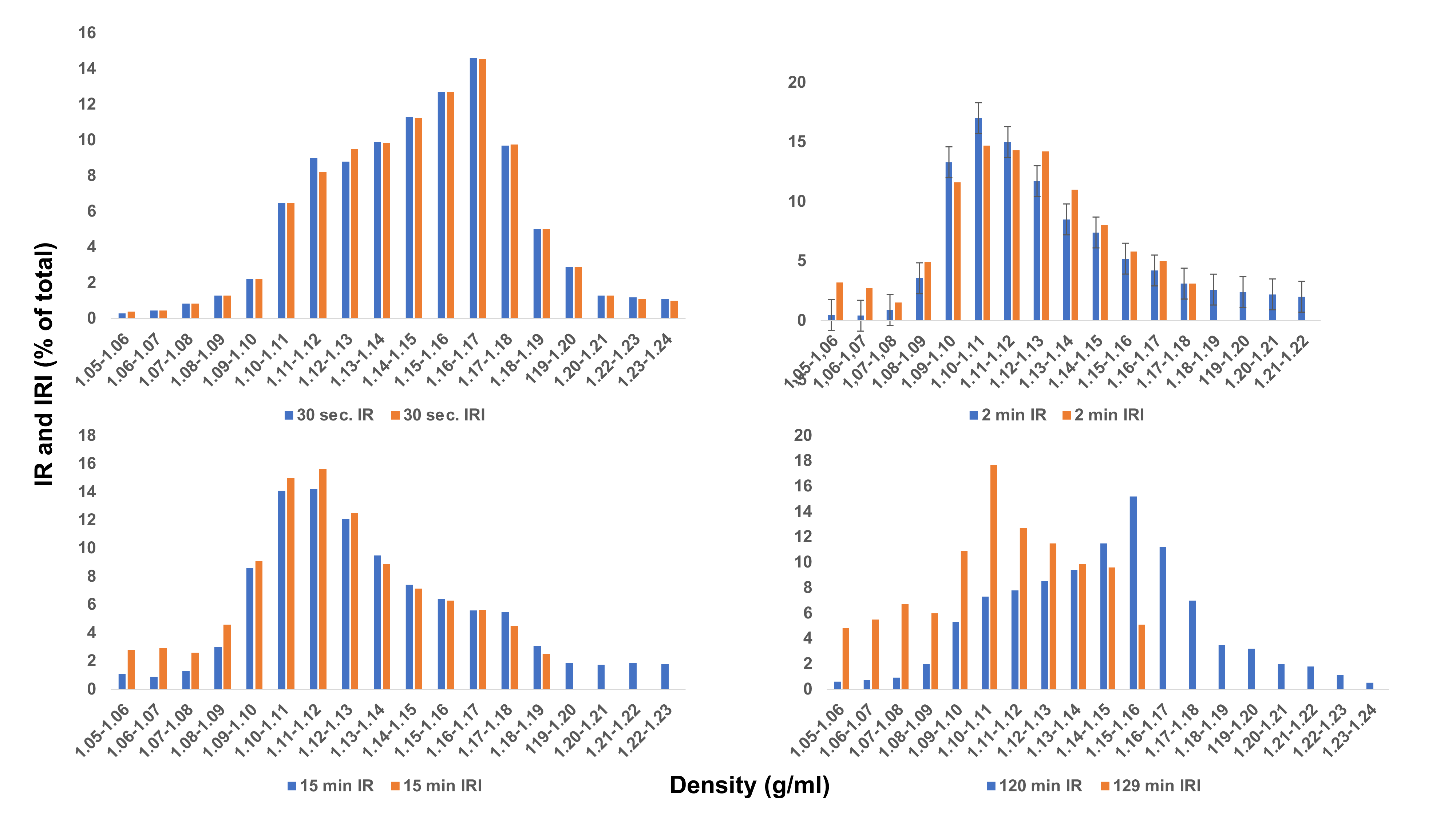
Distribution on density gradients of insulin binding activity and insulin in liver microsomal fractions of control and insulin-treated rats. Microsomal fractions were isolated from rats killed at the indicated time after injection of native insulin (50 μg) or left untreated. Following centrifugation on sucrose density gradients, subfractions were collected and assayed for density, *in vitro* insulin binding activity and extractable immunoreactive insulin content, with results normalized to density intervals of 0.01 g/ml. The results shown are representative of at least three experiments. The positions of galactosyltransferase, 5’nucleotidase and alkaline phosphodiesterase peaks on the gradients (not shown) are the same as in Figure 1.

**Table 1.** Median densities for distribution of protein, marker enzymes, [^125^I]-labeled and native insulin, and insulin binding activity in the microsomal fraction. Liver microsomal fractions isolated from untreated rats and from animals sacrificed after injection of [^125^I]-labeled and native insulin were subjected to sucrose density gradient centrifugation as described in Experimental Procedures. When indicated microsomal fractions were treated by digitonin. About 20 subfractions were collected and assayed for the components indicated. Median densities, expressed as g/ml (mean ± SEM of the indicated number of determinations) were calculated from distributions as a function of density.

**Protein, insulin binding activity and marker enzymes**
|  | Untreated | Digitonin-treated |
| --- | --- | --- |
| Protein | 1.164 ± 0.002 (6) | |
| *In vitro* insulin binding | 1.166 ± 0.002 (5) | 1.181 ± 0.003 ( |
| 5'Nucleotidase | 1.156 ± 0.003 (3) | 1.184 ± 0.002 (4) |
| Alkaline phosphodiesterase | 1.168 ± 0.002 (2) | 1.189 ± 0.002 (4) |
| Glucose 6-phosphatase | 1.206 |  |
| Acid phosphatase | 1.177 ± 0.006 (3) | |
| N-acetyl-β-D-glucosaminidase | 1.200 <u>± 0.005</u> | 1.142 ± 0.002 (3) |
| Galactosyltransferase | 1.125 ± 0.001(3) | 1.153 ± 0.006 (4) |
| IGF2/M6P receptor | 1.127 ± 0.002 (7) | |

***In vivo* [ $^{125}\text{I}$ ]-insulin uptake**
| Time | Untreated* | Untreated | Digitonin-treated |
| --- | --- | --- | --- |
| 30 sec | 1.176 ± 0.002 (5) | 1.164 ± 0.001 (2) | 1.187 ± 0.001 (4) |
| 90 sec | 1.133 ± 0.002 (4) | 1.132 ± 0.006 (2) | 1.179 ± 0.004 (4) |
| 4 min | 1.123 ± 0.004 (7) | 1.120 ± 0.002 (2) | 1.178 ± 0.001 (4) |
| 8 min | 1.117 ± 0.004 ( ) | 1.116 ± 0.004 (2) | 1.171 ± 0.003 (4) |
| 16 min |  | 1.114 |  |
| 24 min |  | 1.112 |  |

***In vitro* insulin binding activity and insulin content after insulin injection**
| Time after injection | Insulin binding activity | Insulin content |
| --- | --- | --- |
| 0 | 1.155 ± 0.007 (6) | not determined |
| 30 sec | $1.147 \pm 0.002$ (2) | $1.146 + 0.003$ (2) |
| 2 min | $1.130 \pm 0.006$ (6) | $1.127 + 0.005$ (4) |
| 5 min | $1.126 \pm 0.008$ (3) | $1.124 + 0.001$ (2) |
| 15 min | $1.118 \pm 0.005$ (6) | $1.118 + 0.002$ (4) |
| 30 min | $1.116 \pm 0.001$ (2) | 1.110 |
| 60 min | $1.135 \pm 0.008$ (5) | $1.114 + 0.006$ (3) |
| 120 min | $1.141 \pm 0.002$ (2) | $1.109 + 0.002$ (2) |

\*In this series the microsomal fraction was layered below the density gradient

The time-dependent changes in the density distribution of acid-extractable insulin following insulin treatment paralleled those of insulin binding activity, but unlike the latter were not reversed after 30 min (Fig 5). Fractionation of light microsomal components (density < 1.15 g/ml) on self-generated Percoll density gradients showed that, up to 60 min after injection, insulin binding activity was recovered at low density, as was galactosyltransferase (Fig. 6). In contrast, the distribution of immunoreactive insulin on Percoll gradients was time-dependently shifted from low to high density, overlapping at late time the position of the lysosomal marker N-acetyl-β-D-glucosaminidase. Similar time-dependent changes in the distribution of [^125^I]-insulin in Golgi/endosome fractions subjected to Percoll density gradients have been observed in untreated and chloroquine-treated rats (Khan *et al*, 1982, 1985). In addition, insulin receptors in lysosomal fractions have been localized on Percoll gradients in a vesicle of intermediate density between Golgi/endosomes and lysosomes (Khan *et al*, 1981).

**Figure 6.**
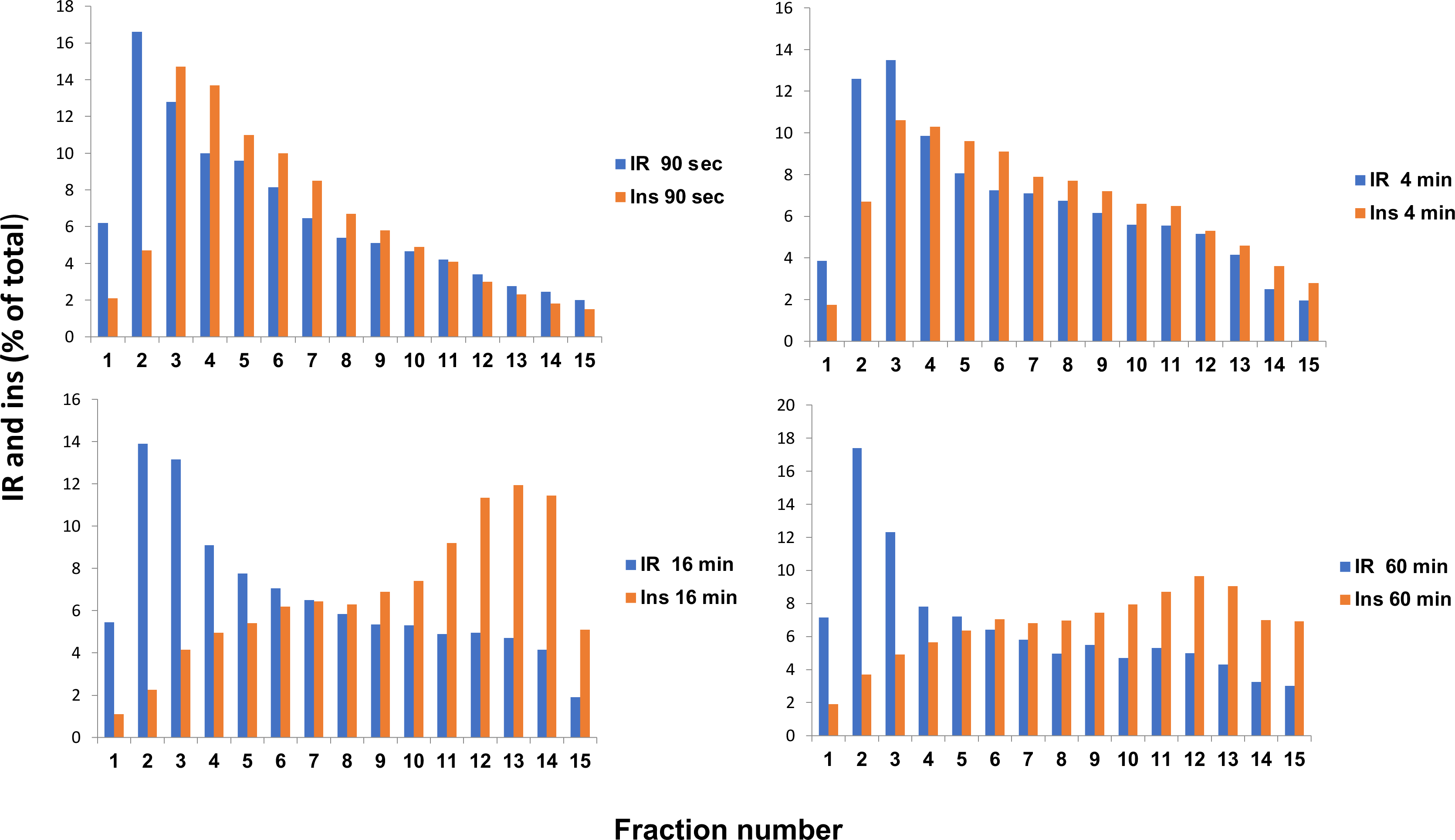
Distribution on Percoll density gradients of insulin binding activity and insulin in liver microsomal subfractions of control and insulin-treated rats. Microsomal subfractions (density < 1.15 g/ml) of rats killed at the indicated time after insulin injection were subjected to centrifugation on Percoll density gradients. Fifteen subfractions of increasing density were collected and assayed for insulin binding activity and extractable immunoreactive insulin content. The results shown are representative of three experiments. Galactosyl transferase was recovered mainly at low density (fractions 1-5) and acid phosphatase at high density (fractions 8-15).

The reversal of insulin-induced changes in insulin receptor distribution, along with the lack of change of total receptor number, suggests that upon acute ligand-induced endocytosis, internalized insulin receptors are fully recycled. However, these results do not exclude the possibility that the late restoration of receptor expression at the cell surface involves in part the synthesis of new receptors. To address this issue, the insulin-induced changes in receptor distribution on sucrose density gradients were examined for sensitivity to *in vivo* treatment with cycloheximide, a protein synthesis inhibitor shown to inhibit insulin receptor synthesis in cultured cells (Knutson *et al*, 1985). As shown on Fig. 7, although *in vitro* insulin binding activity in the microsomal fraction equilibrated at lower densities in cycloheximide treated rats than in control animals, the early insulin-induced shift of activity towards lower densities (at 8 min) and its late reversal (at 3 h) were similar in both groups of rats. Cycloheximide treatment also affected neither the early insulin-induced decrease in insulin binding activity nor its late reversal in the plasma membrane. Finally, the time-dependent translocation of [^125^I]-insulin in the microsomal fraction from high to low density was similar in cycloheximide-treated and untreated rats. Taken together, these observations suggest that following acute insulin-induced endocytosis, internalized insulin receptors are mainly if not fully recycled to the cell surface. In addition, they are consistent with the lack of effect on cycloheximide on the recycling of internalized photoaffinity radiolabeled insulin receptors in hepatocytes (Carpentier *et al*, 1986). As cycloheximide treatment, colchicine treatment of rats did not affect ligand-induced receptor endocytosis and little affected receptor recycling (Postel-Vinay *et al*, 1987).

**Figure 7.**
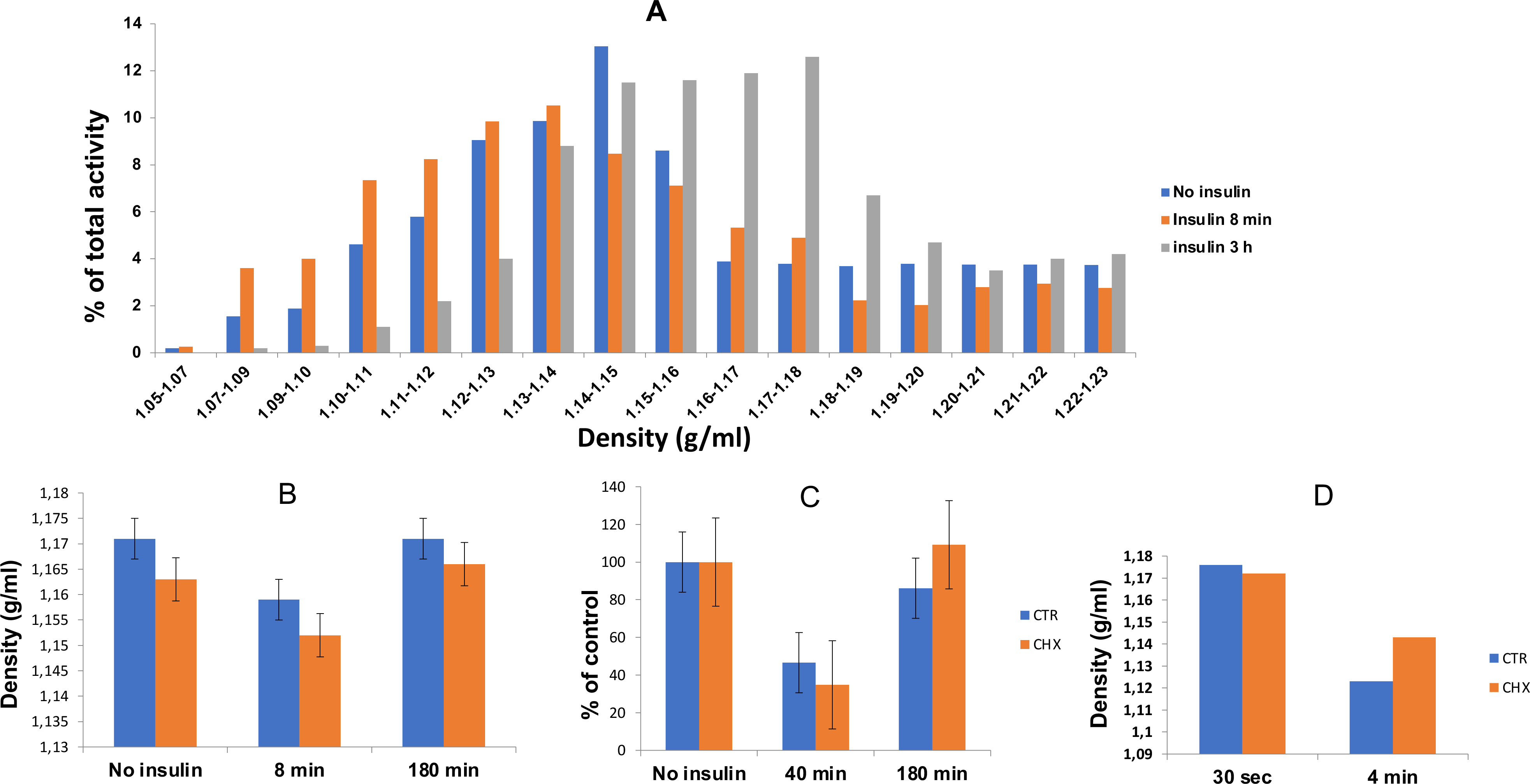
Effect of *in vivo* cycloheximide treatment on the subcellular distribution of insulin binding activity in microsomal fractions of control and insulin-treated rats and of [^125^I]-insulin taken up into liver *in vivo.* A, distribution on sucrose density gradients of insulin binding activity in microsomal fractions of control and insulin-injected rats (8 min and 3 h after injection of 0.3 mg insulin). B, median density of *in vitro* insulin binding activity at the indicated time after insulin injection. C, insulin binding activity in plasma membrane fractions of control and insulin-injected rats (300 μg) at the indicated time after injection. D, median densities of radioactivity in microsomal fractions isolated 30 sec and 4 min in after injection of [^125^I]-insulin. The results shown are representative or the mean of three experiments.

### Physical characterization of the insulin receptor in liver subcellular fractions

The observations described above suggest that, upon ligand-induced endocytosis, the insulin receptor retains functional and presumably structural integrity. To confirm this, the α and β subunits of the receptor in subcellular fractions of control and insulin-injected rats have been characterized by SDS-PAGE in association with affinity crosslinking (α subunit) and Western immunoblotting (both subunits).

As shown in many previous reports, a major insulin binding protein of about 130 kDa (expected Mw for the α subunit) was identified by affinity crosslinking in plasma membrane and Golgi/endosome fractions incubated with [^125^I]-insulin, and was also detectable, albeit to a lesser abundance, in lysosomal fractions (supplementary Fig. 3). An insulin binding protein of 120 kDa, previously identified as a proteolytic product of the α subunit generated *in vitro* but not *in vivo* (Lipson *et al*, 1986, 1988, Sanchez-Casas *et al*, 1995) was also detectable in plasma membranes. As shown previously, the generation of this 120 kDa protein did not occur at 4° C and was inhibited by PMSF, leupeptin and benzamidine.

When the centrifugation step after crosslinking was omitted, two additional proteins, of about 70 and 85 kDa, were detected; the 85 kDa protein bound specifically [^125^I]-insulin, albeit with low affinity (supplementary Fig. 3) To address the possibility that these proteins were proteolytic fragments of the α subunit generated upon receptor endocytosis, subcellular fractions isolated shortly after *in vivo* injection of [^125^I]-insulin were subjected to affinity crosslinking and SDS PAGE analysis (supplementary Fig. 4). As expected from the rapid degradation of internalized [^125^I]-insulin, the expression of the crosslinked 130 kDa protein time-dependently decreased, whereas that of the 70 kDa protein concurrently increased. Further characterization of these proteins after Sephacryl S-300 gel filtration of solubilized endosomal extracts showed that, under nonreducing conditions, the 70 kDa protein was resolved into two components of about 60-65 and 50-55 kDa, respectively.

Although a 55 kDa proteolytic fragment of the insulin receptor has been shown to contain the major crosslinked binding site (Waugh *et al*, 1989), the 70 kDa protein identified under reducing conditions was probably unrelated to the 130 kDa protein (α subunit). Indeed, unlike the latter, it was not immunoprecipitated by two insulin receptor antibodies, did not bind to wheat germ agglutinin and showed no change in electrophoretic mobility following N-glycosidase treatment (results not shown). In addition, its molecular size differed from that of cathepsin D, a 45 kDa soluble aspartic acid protease involved in early steps of insulin degradation in liver endosomes (Authier *et al*, 2002). On the other hand, the molecular size of this protein was closer to that of Hsp 72/73, a heart shock protein identified in cultured fetal hepatocytes (Zachayus *et al*, 1996). The latter protein has been shown to associate with the insulin receptor on co-immunoprecipitation and with [^125^I]-insulin on affinity labeling.

On Western immunoblotting using an antibody against the C-terminal sequence 1336-1355 of the β subunit, a single or major protein of 95 kDa (expected molecular mass for the β subunit) was identified in plasma membrane, Golgi/endosome and lysosome fractions, confirming previous observations (supplementary Fig. 5). As anticipated, on Sephacryl gel filtration of solubilized fractions this 95 kDa protein was co-eluted with insulin binding activity. Two additional minor proteins of lower molecular mass (70-75 and 50-55 kDa), presumably proteolytic fragments of the β subunit, were also detectable in lysosomal fractions, with highest expression in the absence of protease inhibitors.

On Western immunoblotting using an antibody against the 542-551 sequence of the α subunit, two proteins of about 110-115 and 120-125 kDa were identified in the Golgi/endosome fraction and one protein of 140-150 kDa in the plasma membrane fraction (supplementary Fig. 6). As the 95 kDa protein (receptor β subunit), these proteins were also co-eluted with insulin binding activity on Sephacryl gel filtration. Multiple proteins of lower molecular weight (40-110 kDa) were also detected in plasma membrane and lysosomal fractions. Whether the 110-140 kDa proteins mentioned above are related to the α subunit of the insulin receptor is questionable. Indeed, autoradiographic signals were unaffected by the addition of excess 542-551 peptide (results not shown), and their electrophoretic mobility differed from that of the α subunit identified by affinity crosslinking.

Previous studies (Burgess *et al*, 1992, Desbuquois *et al*, 2008) have shown that, as expected from ligand-induced endocytosis of the insulin receptor, the expression of the 95 kDa protein (β subunit) following insulin injection rapidly decreases at the plasma membrane while increasing in endosomes, with a maximum at 15-30 min. Similar insulin-induced changes in the expression of the 135 kDa protein (α subunit) on affinity crosslinking in liver plasma membranes and endosomes have been reported (Sanchez Casas *et al*, 1995). The present study confirms the insulin-induced changes in the expression of the 95 kDa protein in cell fractions and shows that this protein does not undergo proteolysis during redistribution (supplementary Fig. 5). Unexpectedly, the expression of the 120 and 130 kDa proteins identified with the antibody against the α subunit 542-551 sequence in endosomes decreased instead of increasing after insulin injection (supplementary Fig. 6). Although unlikely, association of insulin to this sequence may have resulted in a loss of its accessibility to antibody.

### Cell-free endosome to lysosome transfer of insulin and its receptor

Based on distribution on Nycodenz density gradients and release from lysosomal fractions by lysosome-disrupting drugs, [^125^I]-insulin taken up by rat liver *in vivo* has been shown to undergo a partial transfer from endosomes to lysosomes in a cell-free system (Chauvet *et al*, 1998). So did, and to a greater extent owing to reduced endosomal degradation, [^125^I]-proinsulin (Desbuquois *et al*, 2003). A partial endosome to lysosome transfer of insulin binding activity and receptor β subunit expression also occurred following injection of native insulin (Chauvet *at al*, 1998) and super-active insulin analogs (Authier *et al*, 2004). In the present study, the cell-free transfer of insulin and its receptor has been reexamined using purified endosomal and lysosomal fractions instead of the crude post-mitochondrial supernatant used previously.

Following injection of [^125^I]-insulin, endosomal fractions were isolated and incubated with lysosomal L1 or L2 fractions and cytosol of untreated rats, the concentration of lysosomal protein being varied relative to that of endosomal protein. The distribution of [^125^I]-insulin on Nycodenz density gradients was then examined and compared to that of the lysosomal marker N-acetyl-β−D-glucosaminidase. At 4°C, at least 90% of [^125^I]-insulin was recovered at the endosomal position (densities 1.08-1.10 g/ml), and less than 5% at the lysosomal position (density 1.12-1.14 g/ml). After incubation for 10 min at 37°C, [^125^I]-insulin was in part transferred from the endosomal to the lysosomal position (Figs 8 and 9) However, as the ratio of lysosomal to endosomal protein decreased, the density at which [^125^I]-insulin co-migrated with N-acetyl-β-D-glucosaminidase also decreased, whereas the amount of [^125^I]-insulin recovered at the lysosomal position increased, with comparable results for L1 (Fig. 8) and L2 (Fig. 9) fractions. These observations suggest the formation of hybrid organelles of intermediate density between that of endosomes and that of lysosomes.

**Figure 8.**
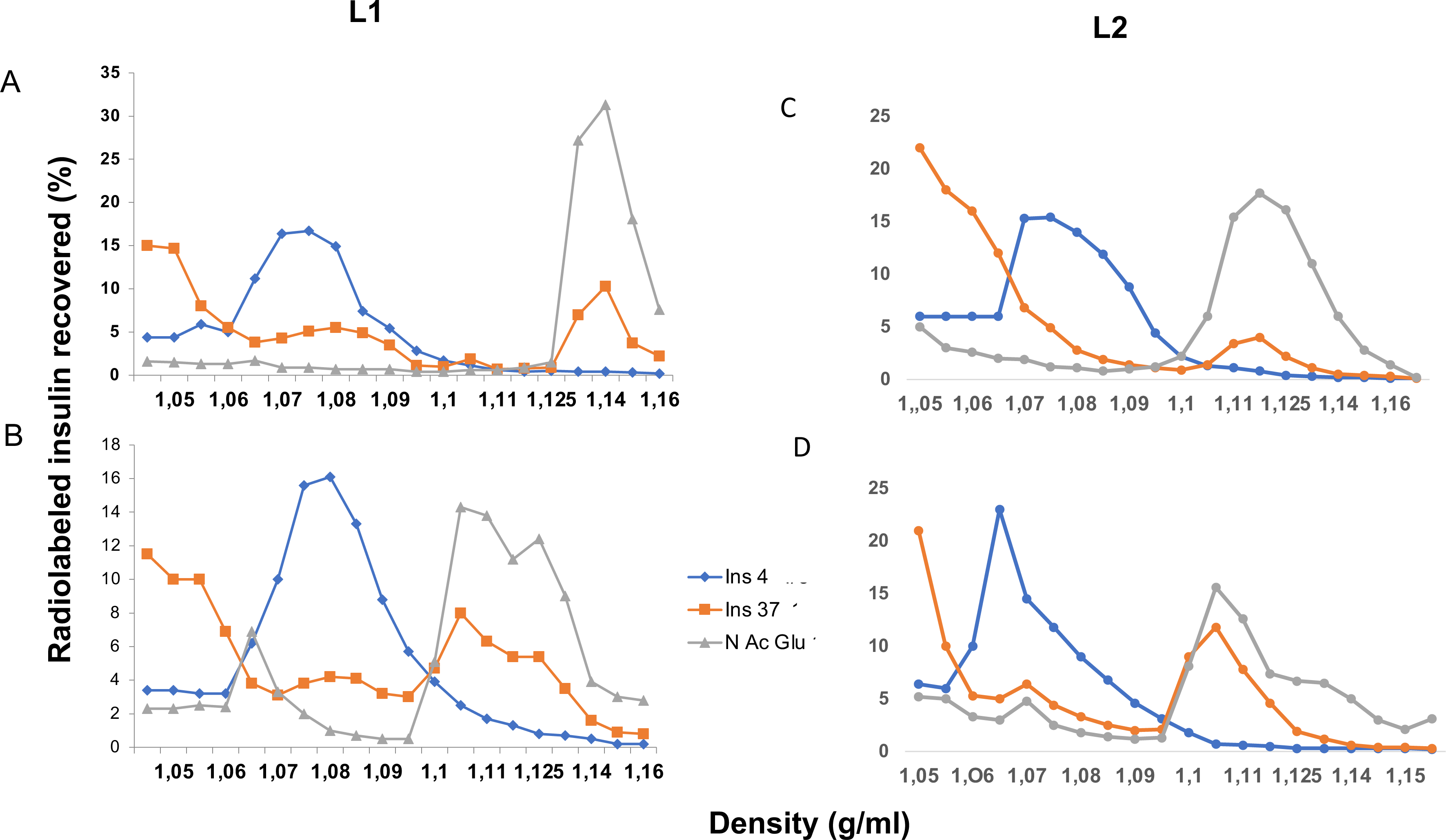
Distribution of [^125^I] insulin taken up into rat liver *in vivo* on density gradients upon cell-free endosome to lysosome transfer. Endosomal fractions isolated 8 min after injection of [^125^I]-insulin were incubated with lysosomal L1 (left) and L2 (right) fractions and cytosol of control rats. The ratios of lysosomal to endosomal protein concentration were, respectively, 6,4 (A) and 0,8 (B) for the L1 fraction, and 7,6 (C) and 1,9 (D) for the L2 fraction. After 10 min at 4°C and 37°C, incubation mixtures were subjected to centrifugation on Nycodenz density gradients. Twenty fractions were collected and assayed for density, radioactivity and N-acetyl-β-D-glucosaminidase activity. Representative distribution profiles for radioactivity and enzyme activity are shown.

**Figure 9.**
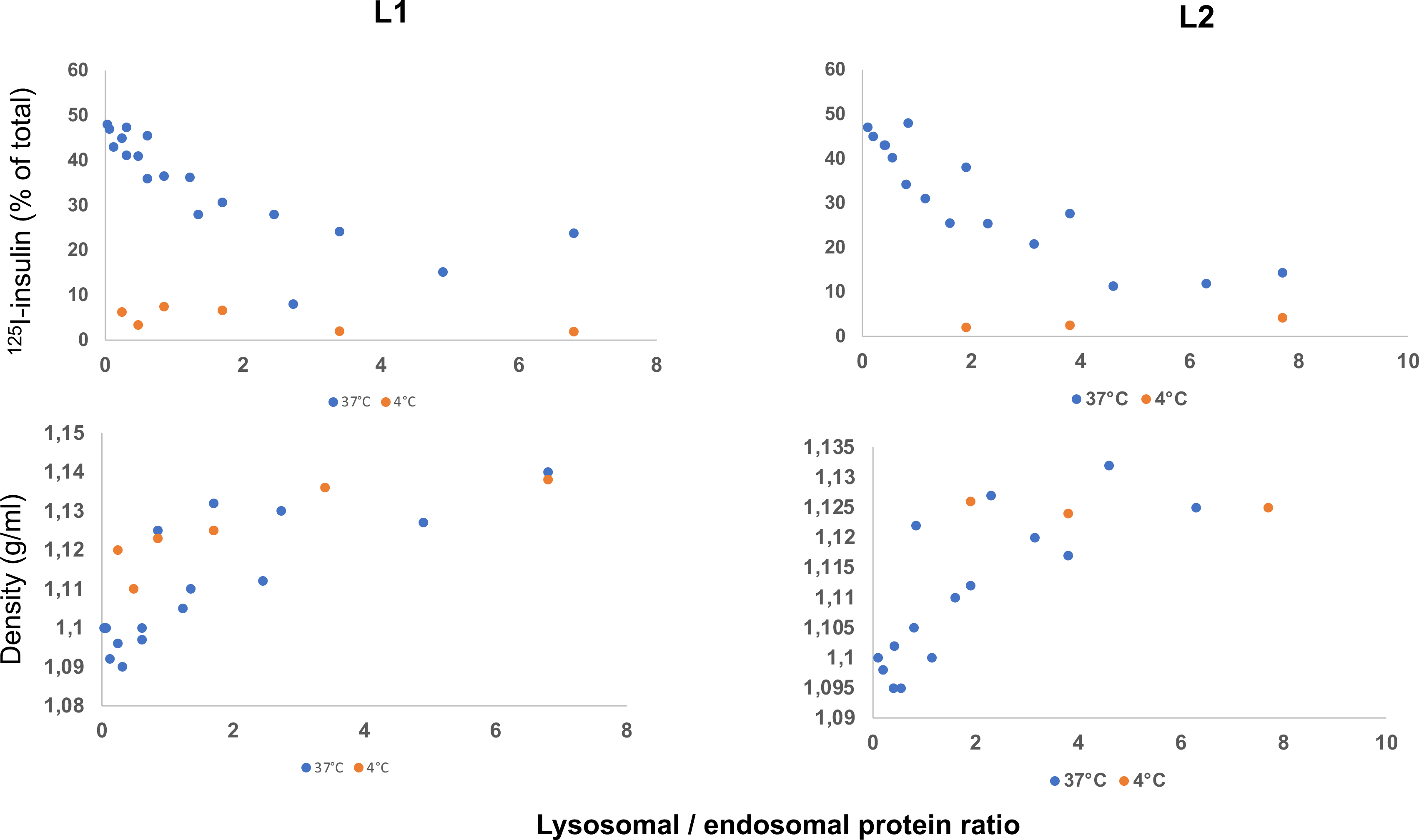
Determination of [^125^I]-insulin migrating at the lysosomal position and of corresponding density as a function of lysosome abundance upon cell-free endosome to lysosome transfer. Endosomal fractions isolated after injection of [^125^I]-insulin were incubated with lysosomal L1 (left) and L2 (right) fractions and cytosol of control rats as described in legend to figure 8. The percentage of [^125^I]-insulin co-migrating with N-acetylβ-glucosaminidase (A) and the corresponding density (B) on Nycodenz density gradients have been plotted against the lysosomal to endosomal protein ratio.

To assess whether the endosome to lysosome transfer of [^125^I]-insulin involves a fusion of these organelles with content mixing, a cell-free assay exploiting the avidin-biotin interaction developed by Mullock *et al* (1994) was used. Endosomes isolated after injection of [^125^I]-biotinylated insulin were incubated with lysosomes isolated after injection of avidin conjugated to galBSA and cytosol, following which the distribution of [^125^I]-biotinylated insulin on Nycodenz density gradients was examined. In a first series of experiments, transfer of [^125^I]-biotinylated insulin to lysosomes isolated after injection of avidin conjugated to galBSA was compared to transfer of [^125^I]-insulin to lysosomes of control rats. Consistent with the *in vivo* studies described above, in this cell-free system [^125^I]-biotinylated insulin was time-dependently transferred from the endosomal to the lysosomal position to a greater extent than [^125^I]-insulin (Fig. 10). In a second series of experiments, the association of [^125^I]-biotinylated insulin with avidin galBSA conjugate during incubation was assessed by immunoprecipitation using anti-avidin antibody. As shown on Fig. 10, while time dependently decreasing in the absence of biocytin or native biotinylated insulin, association of [^125^I]-biotinylated insulin to avidin increased in the presence of these agents. At 10 min, about 3% of total immunoprecipitated [^125^I]-biotinylated insulin (no biocytin present) was bound to avidin in the presence of biocytin.

**Figure 10.**
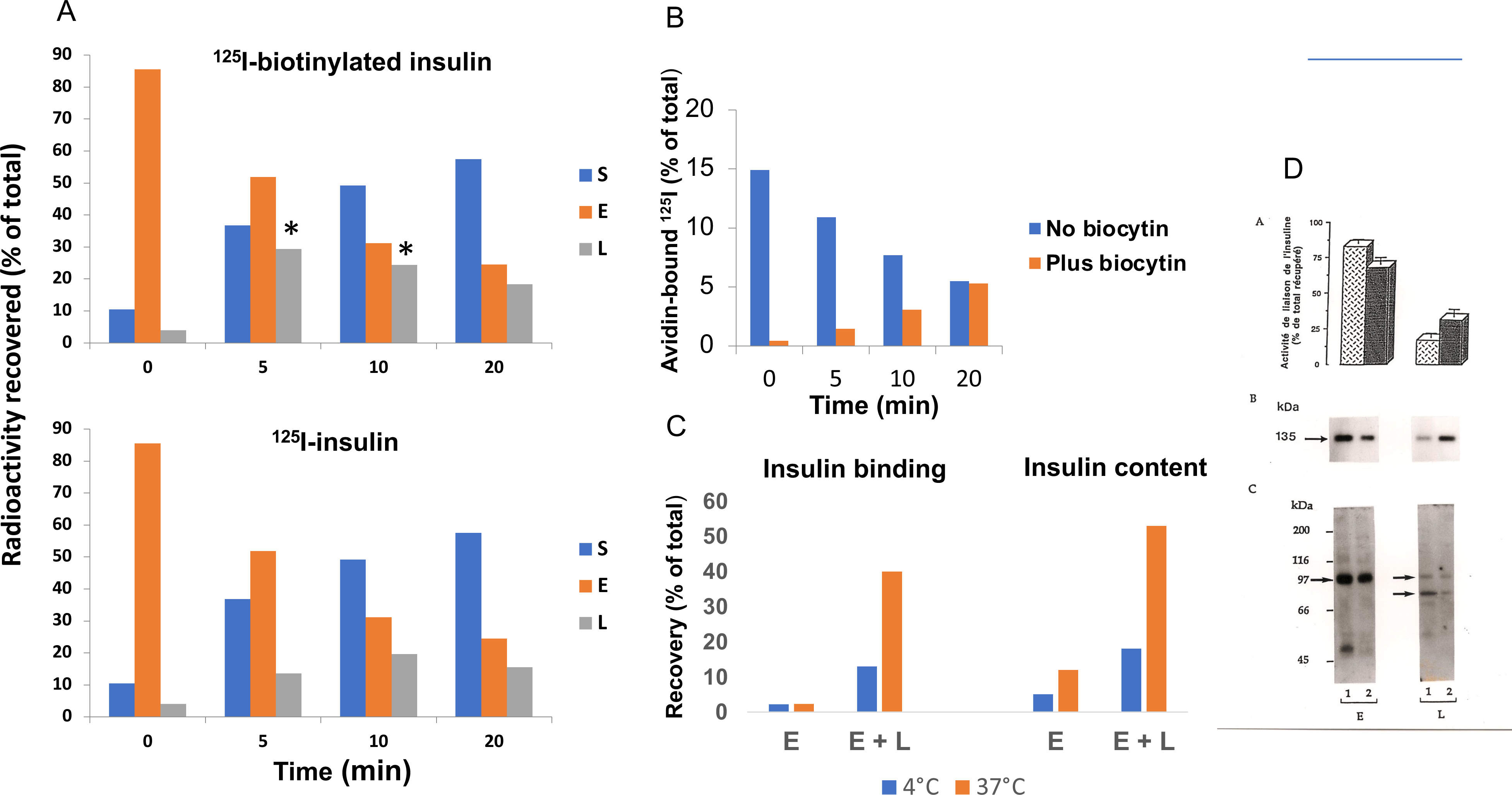
Cell-free endosome to lysosome transfer of [^125^]-biotinylated insulin taken up into rat liver *in vivo*. Liver endosomal fractions of rats isolated 8 min after injection of [^125^I]-biotinylated insulin were mixed with lysosomal (L1 and L2) fractions isolated 30 min after injection of avidin-galBSA conjugate and cytosol. After incubation for 5, 10 and 20 min at 37°C, incubation mixtures were subjected to centrifugation on Nycodenz density gradients and to immunoprecipitation by anti-avidin antibody in the presence and absence of biocytin or native biotinylated insulin. In parallel experiments, endosome fractions of rats killed 8 min after injection of [^125^I]-insulin were incubated with lysosomal and cytosolic fractions of untreated rats prior to density gradient centrifugation. A, comparative distributions of [^125^I]-insulin and [^125^I]-biotinylated insulin in cytosolic, endosomal and lysosomal gradient subfractions (density range, 1.05-1.07, 1.07-1.11 and 1.11-1.15 g/ml, respectively). B, [^125^I]-biotinylated insulin immunoprecipitated by anti-avidin antibody in the absence and presence of biocytin, expressed as % of total in the medium and % immunoprecipitated in the absence of biocytin, respectively. C, cell-free endosome to lysosome transfer of [^125^I]-insulin binding activity (left) and immunoreactive insulin (right). D, cell-free endosome to lysosome transfer of [^125^I]-insulin binding activity (top), insulin receptor α subunit crosslinked to [^125^I]-insulin (middle) and immunodetected β subunit (bottom).

To confirm that the insulin receptor undergoes endosome to lysosome transfer along with insulin under cell-free conditions, endosomal and lysosomal fractions of rats which had been injected with native insulin were mixed and incubated at 37°C in the presence of cytosol. Following centrifugation on Nycodenz gradients, both low density (endosomal) and high density (lysosomal) components were collected and assayed for *in vitro* insulin binding activity and immunoreactive insulin content (Fig. 10). In incubation mixtures left at 4°C, about 14% of insulin binding activity and 18% of immunoreactive insulin were recovered at the lysosomal position. After incubation for 10 min at 37°C, in addition to a 50% decrease of insulin resulting from its endosomal degradation, insulin binding activity and insulin recovered at the lysosomal position were increased by 40% and 55%, respectively. As shown previously (Chauvet *et al*, 1998), in this cell free system an endosome to lysosome transfer of insulin receptor α subunit accompanied the transfer of insulin binding activity (Fig. 10). However, although decreased in the endosomal component, the expression of insulin receptor β subunit did not increase in the lysosomal component, probably as a result of degradation by lysosomal proteases.

## Summary and conclusions

Using density gradients calibrated with markers for cell organelles, we have confirmed that, upon receptor-mediated uptake in rat liver, [^125^I]-insulin undergoes rapid translocation from the plasma membrane to endosomes, with little further translocation to lysosomes. This intracellular pathway differs from that of proteins such as epidermal growth factor and asialoglycoproteins, which upon uptake into liver are in part transferred from endosomes to lysosomes. However, owing to a reduced endosomal degradation, [^125^I]-proinsulin associated to a greater extent than [^125^I]-insulin with lysosomes, as did [^125^I]-biotinylated insulin, the α amino group of which is blocked.

We have also confirmed that, following injection of native insulin, both insulin and its receptor in liver are rapidly translocated from the plasma membrane to endosomes. However, whereas at late times the internalized receptor was recycled to the plasma membrane, internalized insulin time-dependently segregated from the receptor and associated with endosomal components of higher density enriched in acid phosphatase. Neither receptor and insulin endocytosis, nor receptor recycling were affected by cycloheximide, an inhibitor of protein synthesis.

Using affinity crosslinking and Western immunoblotting procedures, we have confirmed that α and β subunits of the insulin receptor in liver subcellular fractions are expressed as proteins of about 135 kDa and 95 kDa, respectively. Following insulin treatment, the β subunit as detected by Western immunoblotting was rapidly translocated from the plasma membranes to endosomes. The 70 kDa insulin binding protein identified by affinity crosslinking in endosomes was probably unrelated to the receptor α subunit.

A time-dependent endosome to lysosome transfer of *in vivo* internalized [^125^I]-insulin and [^125^]-biotinylated insulin occurred in a liver cell-free system. As shown previously for the transfer of [^125^I]-asialofetuin (Mullock *et al*, l989), the transfer of [^125^I]-insulin was function of the abundance of lysosomes relative to that of endosomes. Upon cell-free transfer from endosomes to lysosomes loaded with avidin galBSA conjugate, [^125^I]-biotinylated insulin was in part immunoprecipitated by anti-avidin antibody, consistent with endosome to lysosome fusion with organelle content mixing. However, association of [^125^]-biotinylated insulin with avidin galBSA conjugate was lower than reported for association of endosomal avidin-asialofetuin with lysosomal [^125^I]-biotinylated polymeric IgA (Mullock *et al*, 1994, 1998). This may have resulted from a more rapid degradation of [^125^I]-biotinylated insulin, as compared to that of [^125^I]-biotinylated polymeric IgA.

## Experimental procedures

### Materials

Porcine insulin and recombinant human insulin were purchased from Novo Nordisk. Porcine proinsulin, des-alanine des-asparagine insulin and des-octapeptide insulin were gifts from Eli Lilly Research laboratories. Biotinylated insulin, avidin, N-succinimidyl 3-(2-pyridyldithio)propionate (SPDP), protein A, cycloheximide, and antibodies against insulin and avidin were from Sigma-Aldrich. Antibodies against the β subunit of the insulin receptor were from Santa Cruz Biotechnology. Antibodies against partial sequences of α (542-551) and β(1336-1355) subunits of the human insulin receptor were gifts of Drs Emmanuel Van Obberghen and Daniel Lane, respectively. Nitrocellulose membranes were from Biorad. Other chemicals were obtained commercially as described previously.

### Preparation of galBSA and conjugation of galBSA to avidin

GalBSA was prepared from bovine serum-albumin by reductive lactosamination as described by Wilson (1978). Conjugation of galBSA to avidin was performed using SPDP as described by Carllson *et al* (1978). Briefly, 8 mg (0.12 μmol) of galBSA diluted into 1.8 ml of 0.1 M phosphate buffer, pH 7, 0.1 M NaCl, was reacted with 0.3 mg (0.93 μmol) of SPDP dissolved into 30 μl of ethanol. After 30 min at room temperature, the reaction mixture was dialyzed against 500 ml of 0.1 M acetate buffer, pH 4.5, 0.1 M NaCl. Based on measurements of absorbance at 280 and 343 nm after addition of 50 μl of 50 mM dithiothreitol (DTT), the number of SPDP groups introduced into galBSA was estimated as about 7 per molecule. Reaction of avidin (5,4 mg, 0.08 μmol) with SPDP (0,15 mg, 0.46 μmol) was carried out n a similar manner. Following reduction by DTT, thiol-containing gal BSA was coupled to avidin-2-pyridyl disulphide by thiol-disulfide exchange. After 20 h at 23°C, the reaction mixture was subjected to gel filtration on Sephacryl S300. About 70% of total applied protein as measured by absorbance at 280 nm was eluted ahead of BSA (66 kDa), close to the elution position of alcohol dehydrogenase (150 kDa).

### Preparation of [^125^I]-labeled polypeptides

[^125^I]-labeled insulin, biotinylated insulin and proinsulin (400-600 Ci/mmol) were prepared using lactoperoxidase and purified by gel-filtration on Sephadex G50. As shown in a previous study (Desbuquois *et al*, 2003), iodination of insulin occurred mainly (70%) at tyrosine A14. [^125^I]-labeled protein A was prepared and purified in a similar manner.

### Animal experiments

*In vivo* procedures were approved by the French institutional committee for use and care of experimental animals. Male Sprague Dawley rats (body weight, 200 g), obtained from Charles River France and Elevage Janvier, were fasted for 16 h prior to experiments. Native insulin, insulin derivatives and proinsulin, [^125^I| labeled insulin and biotinylated insulin (2-5 pmol each) and avidin conjugated to galBSA were diluted into 0.5 ml of physiological saline and injected into the penis vein under ether or Nembutal anesthesia. When indicated rats injected with native and [^125^I]-labeled insulin received an intraperitoneal injection of cycloheximide (0.8 mg per 100 g of body weight) 3 h prior to injection. At the indicated times, the liver was removed and immediately homogenized in 0.25 M sucrose.

### Liver subcellular fractionation

This was performed using preparative and analytical and preparative procedures as described previously (Desbuquois *et al*, 1979, Posner *et al*, 1980, Bergeron *et al*, 1986). Nuclear (N), mitochondrial-lysosomal (ML), light mitochondrial (L) and microsomal (P) fractions were prepared from liver homogenates by differential centrifugation. In analytical procedures, these primary fractions were quantitatively further fractionated by centrifugation on linear sucrose (N, ML and P fractions) and Nycodenz (L fraction) density gradients. When indicated the microsomal fraction was treated by digitonin (0.17%) for 30 min prior to density gradient subfractionation. Sucrose gradients (density range, 1.04-1.25 g/ml) were centrifuged for 15 h at 27,000 rpm in a Beckman SW 28 rotor, and Nycodenz gradients (density range, 1.04-1.16 g/ml) for 1 h at 30.000 rpm in a Beckman SW41 rotor. About 15-20 fractions were collected and assayed for density by refractometry. Low density (d < 1.15 g/ml) microsomal subfractions were further fractionated in a Percoll density gradient (32% Percoll in 0.25 M sucrose) generated by centrifugation for 30 min at 30,000 rpm in a Beckman 50Ti rotor. Purified plasma membranes were isolated from the N fraction according to Hubbard *et al* (1983), Golgi-endosomal fractions from the P fraction according to Ehrenreich *et al* (1973), and lysosomal L1 and L2 fractions from the L fraction according to Wattiaux *et al* (1978).

Subcellular fractions isolated after injection of [^125^I]-labeled insulin, biotinylated insulin and proinsulin were analyzed for radioactivity, and fractions isolated from control and insulin-injected rats for *in vitro* insulin binding activity and acid-extractable insulin content as described previously (Desbuquois *et al*, 1982, 2003). The distribution of these components on sucrose and Nycodenz density gradients was compared to that of markers for plasma membranes (5’nucleotidase and alkaline phosphodiesterase), Golgi (galactosyltransferase), endosomes (IGF2/mannose 6-phosphate receptor) and lysosomes (acid phosphatase and N-acetyl β-D-glucosaminidase). Median densities for enzymes, [^125^I]-insulin, *in vitro* insulin binding activity and immunoreactive insulin in the P fraction are shown in Table 1, and representative distributions of several enzymes and insulin binding activity in supplementary Fig. 8.

### Physical characterization of insulin receptor subunits in liver cell fractions

The α subunit of the insulin receptor was characterized by cross-linking to *in vitro* added or *in vivo* injected [^125^I]-insulin with DSS followed by SDS PAGE and autoradiography. Both subunits were also analyzed by Western immunoblotting with antibodies against sequences 542-551 (α subunit) and 1336-1355 (β subunit) of the human insulin receptor. When indicated insulin receptor crosslinked to [^125^I]-insulin was characterized after gel filtration of Triton X-100 solubilized fractions on Sephacryl S-300. [^125^I]-labeled protein A was used to reveal immune complexes in Western blots. In some experiments, a commercial goat polyclonal antibody against insulin receptor β subunit was used and immune complexes were revealed by enhanced chemiluminescence detection.

### Endosome to lysosome transfer of *in vivo* internalized [^125^I]-insulin, [^125^I]-biotinylated insulin, native insulin and insulin receptor in a cell-free system

Liver Golgi/endosome fractions of rats sacrificed 8 min after injection of [^125^I]-insulin were incubated with lysosomal and cytosolic fractions of control rats in 0.5-1 ml of 0.25 M sucrose/5 mM TES containing 1 mM ATP, 5 mM MgCl2, 5 mM phosphocreatine and 0.5 mg/ml creatine kinase. After 5-20 min at 37°C, incubation mixtures were cooled to 4°C and centrifuged on linear Nycodenz gradients (density range, 1.04-1.16 g/ml) calibrated with the lysosomal enzyme N-acetyl β-D-glucosaminidase. Endosome to lysosome transfer of [^125^I]-biotinylated insulin was studied in a similar manner but in this case lysosomal fractions were isolated 30 min after injection of avidin conjugated to galBSA. Association of [^125^I]-biotinylated insulin loaded endosomes with avidin-galBSA loaded lysosomes was assessed using a content mixing assay as described by Mullock *et al* (1994). Briefly, after incubation in presence of native biotinylated insulin (10 μg/ml) or biocytin (0,2 mg/ml), incubation mixtures were diluted into 50 mM Tris buffer, pH 7.5, containing 1% Triton X-100 and protease inhibitors (PMSF 1 mM, benzamidine l mM, bacitracin 1 mg/m1, 1,10 phenanthroline 5 mM, pepstatin 1 μM). Following incubation of the lysed mixtures with anti-avidin antibody at 4°C for 24 h, protein A-Sepharose was added and immunoprecipitated [^125^I]-biotinylated insulin was determined. It was expressed as % of the total mmunoprecipitated in the absence of biotinyl blocking agents.

### Measurement of insulin concentration and insulin binding activity in liver subcellular fractions

The concentration of insulin in acid extracts of liver subcellular fractions was measured by radioimmunoassay using [^125^I]-insulin and anti-insulin antibody. Insulin binding activity in Triton X-100 solubilized subcellular fractions was measured using [^125^I]-insulin as ligand and polyethyleneglycol to precipitate insulin-receptor complexes. Both assays have been described previously (Desbuquois *et al*, 1982, 2003).

## Author contribution

Khadija Tahiri has been involved in the physical characterization of the insulin receptor in liver subcellular fractions; Françoise Fouque, in the biochemical characterization of these fractions; and Marie-Catherine Postel-Vinay, in the study of the effect of cycloheximide on [^125^I]-insulin and insulin receptor trafficking. Bernard Desbuquois coordinated these studies and wrote the manuscript.

## Aknowledgments

The authors gratefully acknowledge the technical assistance of Henriette Burlet, Brigitte de Gallé, Aline Dupuis and Christine Kayser.

This work is dedicated to the memory of Geneviève Chauvet, who initiated the studies on the cell-free endosome to lysosome transfer of insulin and its receptor.

### Abbreviations

Akt: Akt serine/threonine kinase
APS: adaptor protein with PH and SH2 domains
CEACAM: carcino-embryonic antigen-related cell adhesion molecule
DSS: disuccinimidyl suberate
galBSA: galactosylated bovine serum-albumin
Grb: growth factor receptor bound protein
IRS: insulin receptor substrate
PC-1/ENPP1: plasma cell membrane glycoprotein 1/ectonucleotide pyrophosphatase phosphodiesterase 1
PI3K: phosphoinositide-3-kinase
Shc: Src homologous and collagen protein
SOCS: suppressor of cytokine signaling
SPDP: N-succinimidyl 3-(2-pyridyldithio) propionate
WGA: wheat germ agglutinin.

## Supplementary figures legends

**Supplementary Fig. 1. Dose dependent inhibition of [^125^I]-insulin binding to liver microsomal membranes by native insulin and related polypeptides**. Native insulin, des-alanine des-asparagine insulin, des-octapeptide insulin and proinsulin were examined at the indicated concentrations for their ability to inhibit the *in vivo* uptake of [^125^I]-insulin into liver microsomal fractions.

**Supplementary Fig. 2. In vivo uptake of [^125^I]-insulin by rat tissues.** [^125^I]-insulin, alone and with excess native insulin (50 μg), was injected to rats and examined for its ability to be taken up by the particulate fraction of the indicated tissues at 2 min post injection. Nonspecific uptake was determined after co-injection of [^125^I]-inulin and excess native insulin.

**Supplementary Fig. 3. SDS PAGE analysis of proteins crosslinked to *in vitro* added [^125^I]-insulin**. A, proteins identified in microsomal (P), plasma membrane (MP), Golgi/endosomal (GE) and lysosomal (L1 and L2) fractions. B, proteins identified in the plasma membrane fraction incubated with [^125^I]-insulin for 10 min (1), 30 min (2), 60 min (3) and 120 min (4) in the absence of protease inhibitors (left), and for 2 h in the presence of 1 mM PMSF (a), 2 mg/ml bacitracin (b), 10 mg/ml leupeptin (c), 5 mM 1,10 phenantroline (d), 1 mM benzamidine (e) and 1 mM N-ethylmaleimide (f) (right). C, proteins identified in plasma membrane (MP) and Golgi/endosomal (GE) fractions incubated with [^125^I]-insulin for 60 min at 4° C (a), and for 5 min (b), 15 min (c) and 1 h (d) at 30° C. D, [^125^I]-labeled proteins identified in cell fractions that were not subjected to centrifugation. Left, proteins in Golgi/endosomal (GE) and lysosomal (L1 and L2) fractions in the absence (1) and presence (2) of excess native insulin. Right: proteins in the GE fraction in the absence (1) and presence of native insulin at concentrations of 0,04 (2), 0,2 (3), 1 (4) and 5 (5) μg/ml.

**Supplementary Fig. 4. SDS PAGE analysis of proteins crosslinked to *in vivo* injected [^125^I] insulin**. A, proteins identified under reducing conditions in plasma membrane (MP) and Golgi/endosomal (GE) fractions isolated 2 min (a), 5 min (b) and 10 min (c) after injection of [^125^I] insulin (left), and in the Golgi/endosomal (GE) fraction isolated at 10 min in the absence (c) and presence of injected chloroquine (d) and bacitracin (e) (right). Arrows indicate the position of the 135 kDa and 70 kDa [^125^I]-labeled proteins. The determination of the radioactivity associated with these two proteins and of the total TCA-soluble radioactivity as a function of time is shown below. B, chromatographic elution profile and SDS PAGE analysis of endosomal proteins crosslinked to *in vivo* injected [^125^I]-insulin. Proteins of a Golgi/endosomal fraction isolated 6 min after injection have been crosslinked to [^125^I]-insulin, solubilized by Triton X-100 (0.33%) and subjected to chromatography on a Sephacryl S-300 column (1 x 50 cms) equilibrated in 0.1 M ammonium acetate buffer, pH 7.4, calibrated with molecular weight markers. Lower figure: elution pattern of the radioactivity showing two components (peak 1, fractions 16-22, and peak 2, fractions 32-35) eluted ahead of [^125^I]-insulin (peak 3, fractions 40-60). Arrows indicate the elution positions of dextran blue (BD), thyroglobulin (TG, 669 kDa), apoferritin (AF, 443 kDa), alcohol dehydrogenase (AD), 150 kDa) and serum-albumin (SA, 66 kDa). Upper figures: SDS-PAGE analysis and autoradiography of eluate fractions showing, under reducing conditions (upper gel) a 135 kDa protein in peak 1 (fractions 19 and 20) and a 70 kDa protein in peak 2 (fractions 30-32), and under non-reducing conditions (lower gel), a protein of > 200 kDa in peak 1 (fractions 24-26) and two proteins, of about 65 and 50 kDa, in peak 2 (fractions 30-34). C, SDS PAGE analysis of ^125^I-labeled proteins identified in peaks 1 and 2 of Sephacryl S-300 chromatographic eluates. Left and middle: proteins identified under nonreducing conditions following adsorption of eluates peaks 1 and 2 on WGA-Sepharose 6MB (lanes 1, precipitates; lanes 2, supernatants). Right: comparative migration profile, under nonreducing conditions, of peak 2 endosomal protein cross-linked to [^125^I]-insulin (lane 1), [^125^I]-labeled serum-albumin (lane 2) and [^125^I]-labeled heat shock protein (hsp) 70 (lane 3).

**Supplementary Fig 5. Western blot analysis of proteins identified in liver subcellular fractions of control and insulin-injected rats using an antibody against insulin receptor βsubunit**. Left: lane 1, microsomal fraction of control rats; lanes 2 and 3, plasma membrane fraction of control (2) and insulin-treated (3) rats; lanes 4 and 5, Golgi/endosome fraction of control (4) and insulin-treated (5) rats; lanes 6-8, lysosomal L (6), L1 (7) and L2 (8) fractions of control rats. Right: plasma membrane (PM) and Golgi/endosomal (GE) fractions of control and insulin-injected rats at the indicated time after insulin injection

**Supplementary Fig. 6. Western blot analysis of proteins identified in liver subcellular fractions of control and insulin-treated rats using an antibody against insulin receptor α subunit.** Left: lane 1, microsomal fraction of control rats; lanes 2-4, plasma membrane fraction of control (2, 3) and insulin-treated (4) rats; lanes 5-7, Golgi/endosomal fraction of control (5) and insulin-treated (6, 7) rats; lanes 8-10, lysosomal L (8), L1 (9) andL2 (10) fractions of control rats. Right: plasma membrane (PM) and Golgi/endosomal (GE) fractions of control and insulin-injected rats at the indicated time after insulin injection.

**Supplementary Fig. 7. Chromatographic elution profile of proteins immunodetected in liver plasma membrane and Golgi/endosome fractions.** Plasma membrane (PM) and Golgi/endosome (GE) fractions have been subjected to gel filtration on a Sephacryl S-300 column (1 x 50 cms) after solubilization by Triton X-100. Initial (T) and eluate fractions (numbered 22-30) have been probed by Western immunoblotting using antibodies against the α (top) and β (bottom) subunits of the insulin receptor. Insulin binding activity as measured using [^125^I]-insulin was recovered in eluate fractions 22-26.

**Supplementary Fig. 8. Density distribution of enzymes and insulin receptor in the liver microsomal fraction.** A microsomal fraction of control rats has been subjected to sucrose density gradient centrifugation, following which the distribution of galactosyltransferase (Gtr), IGF2/mannose 6-phosphate receptor (IG2/M6P), alkaline phosphodiesterase (PDE) and insulin receptor (IR) has been plotted against normalized density intervals of 0.01 g/ml. The results shown are representative of at least three separate experiments.

## Notes

### Competing Interest Statement

There are no competing interests

### Summary of Updates

There are only very minor changes in this revised version. In figure 7, labels on the ordinate and abscissa of the graphs have been added. In the text, the name of Christine Kayser has been added in the akowledgments.

