## Supplemental figures 1-8 for "Revisiting the trafficking of insulin and its receptor in rat liver: *in vivo* and cell-free studies"

### Slide 1
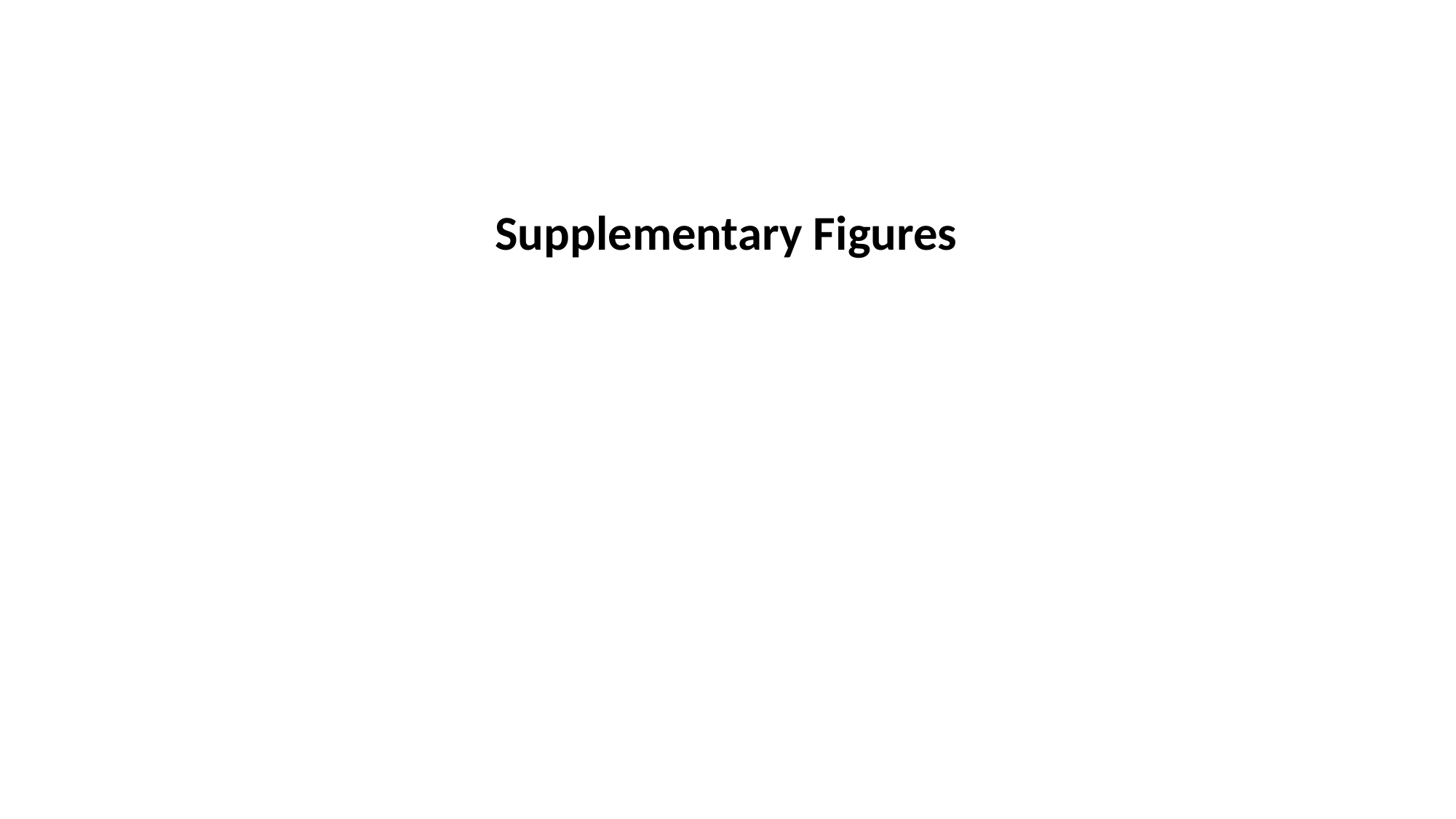

Supplementary Figures

### Slide 2
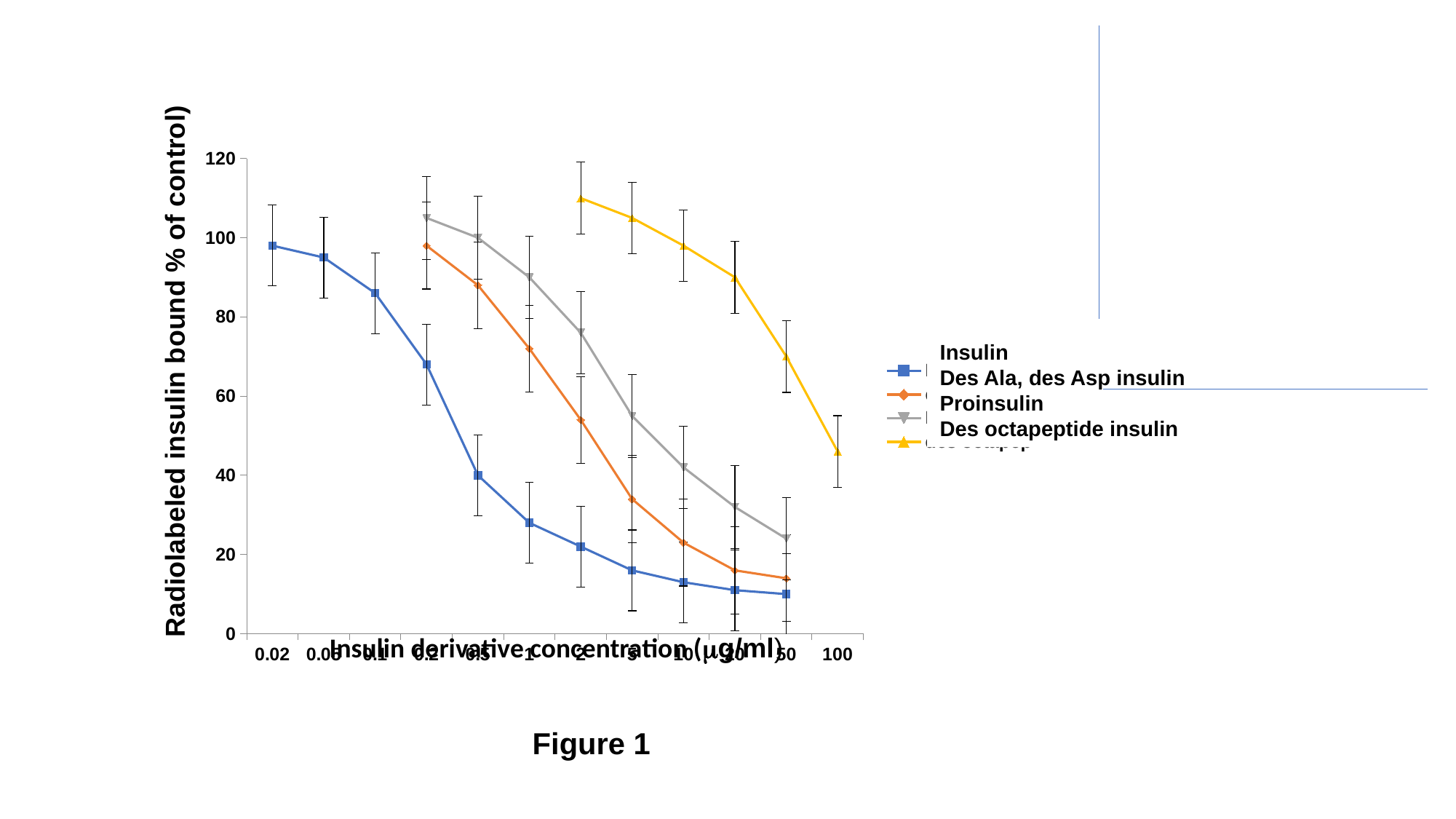

#### Chart
| Category | Insuline | des Ala, Asp | Proinsuline | des octapep |
|---|---|---|---|---|
| 0.02 | 98.0 | None | None | None |
| 0.05 | 95.0 | None | None | None |
| 0.1 | 86.0 | None | None | None |
| 0.2 | 68.0 | 98.0 | 105.0 | None |
| 0.5 | 40.0 | 88.0 | 100.0 | None |
| 1 | 28.0 | 72.0 | 90.0 | None |
| 2 | 22.0 | 54.0 | 76.0 | 110.0 |
| 5 | 16.0 | 34.0 | 55.0 | 105.0 |
| 10 | 13.0 | 23.0 | 42.0 | 98.0 |
| 20 | 11.0 | 16.0 | 32.0 | 90.0 |
| 50 | 10.0 | 14.0 | 24.0 | 70.0 |
| 100 | None | None | None | 46.0 |
Insulin
Des Ala, des Asp insulin
Proinsulin
Des octapeptide insulin
Radiolabeled insulin bound % of control)
Insulin derivative concentration (μg/ml)
Figure 1

### Slide 3
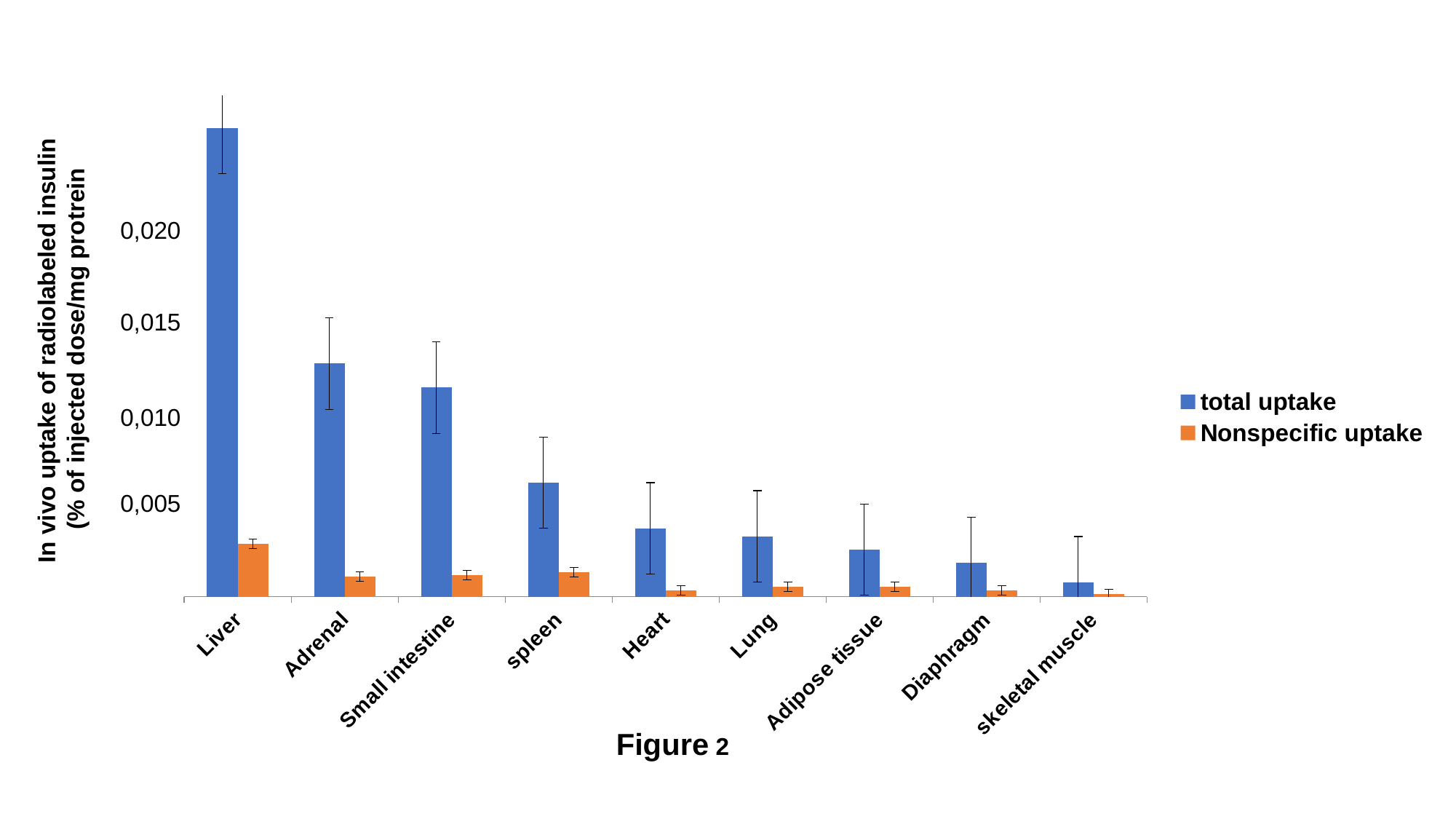

#### Chart
| Category | total uptake | Nonspecific uptake |
|---|---|---|
| Liver | 187.0 | 21.0 |
| Adrenal | 93.0 | 8.0 |
| Small intestine | 83.4 | 8.6 |
| spleen | 45.4 | 9.8 |
| Heart | 27.2 | 2.4 |
| Lung | 24.0 | 4.0 |
| Adipose tissue | 18.7 | 3.9 |
| Diaphragm | 13.5 | 2.5 |
| skeletal muscle | 5.7 | 1.1 |0,020
0,015
In vivo uptake of radiolabeled insulin
 (% of injected dose/mg protrein
0,010
0,005
Figure 2

### Slide 4
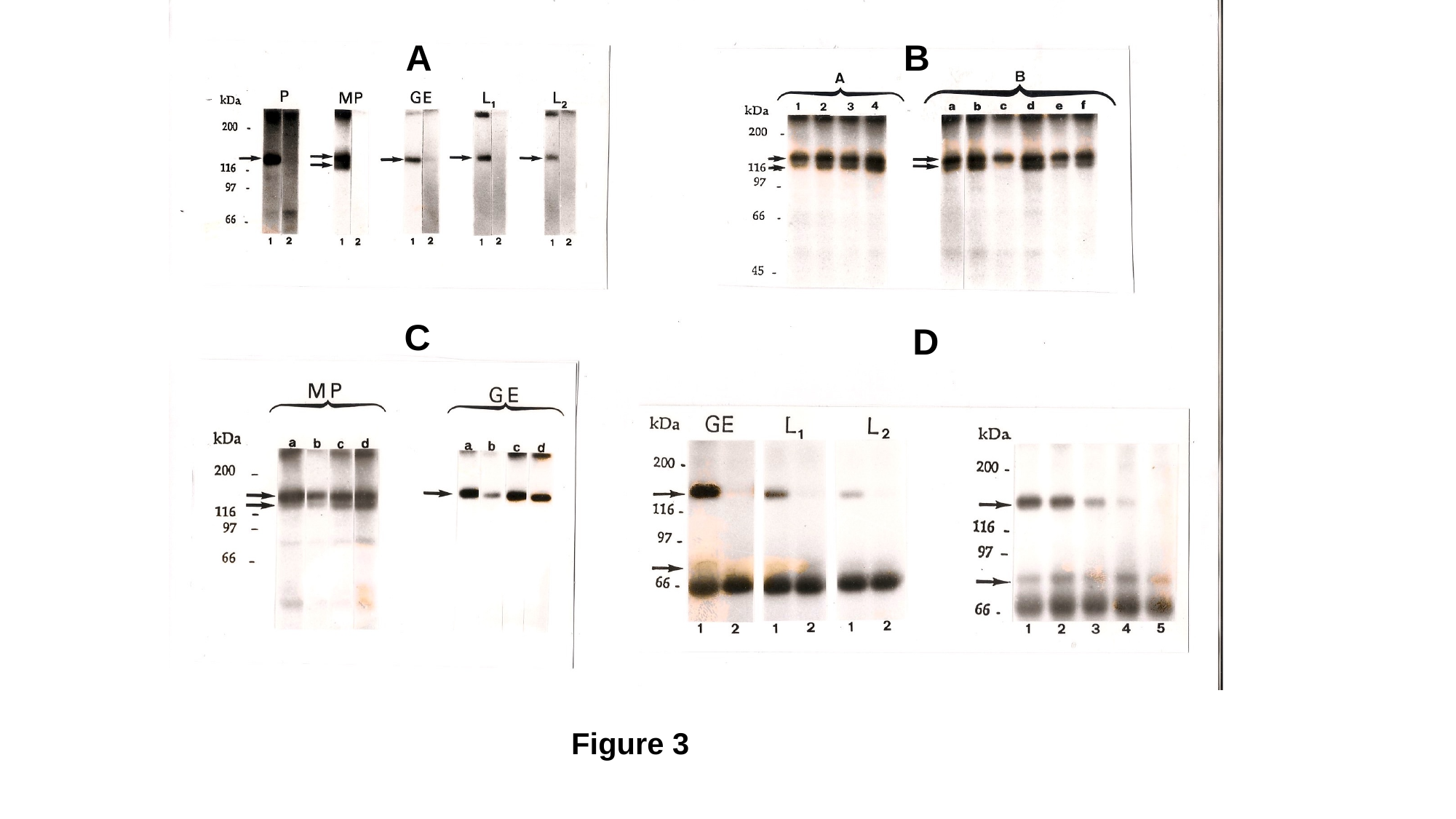

A
B
C
D
Figure 3

### Slide 5
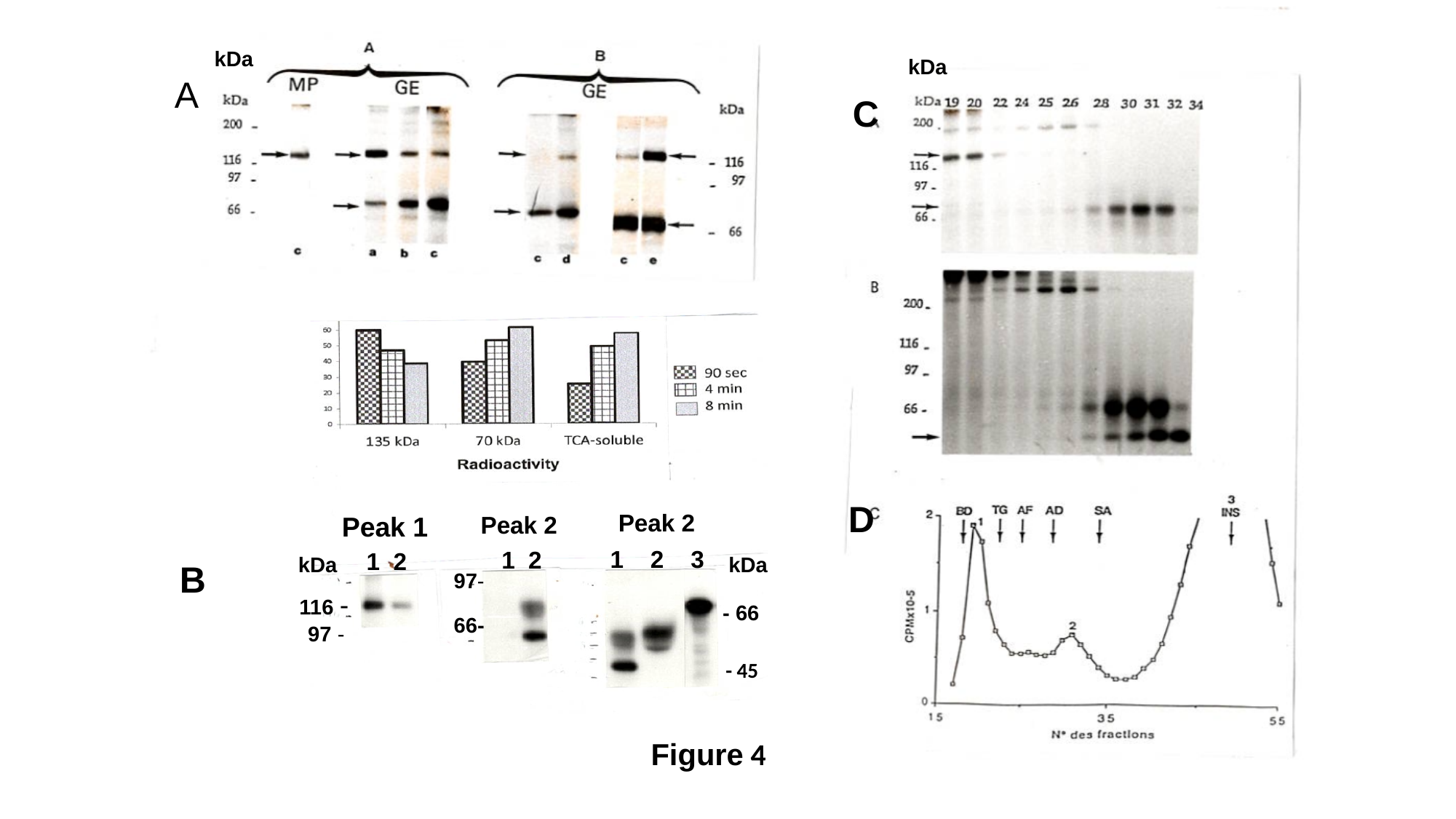

A
C
B
D
kDa
kDa
Peak 2
Peak 1
Peak 2
1 2 3
1 2
1 2
kDa
kDa
97-
200 -
116 -
75 -
97 -
- 66
66-
116 -
97 -
60 -
66 -
- 45
45 -
Figure 4
Figure 4

### Slide 6
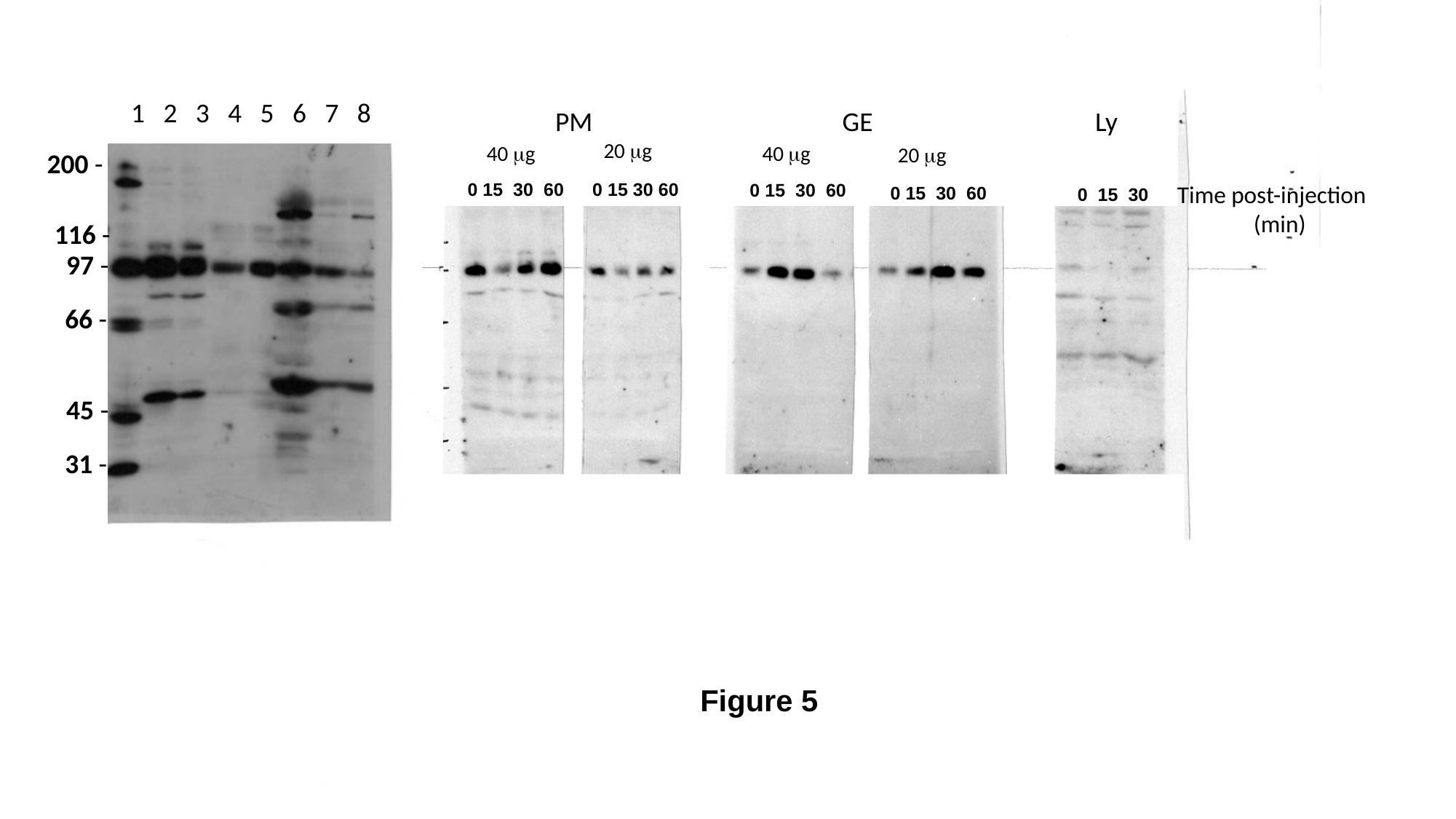

E
1 2 3 4 5 6 7 8
kDa
200 -
116 -
97 -
66 -
45 -
31 -
PM
GE
Ly
20 μg
40 μg
40 μg
20 μg
0 15 30 60
0 15 30 60
0 15 30 60
Time post-injection
 (min)
0 15 30 60
0 15 30
Figure 5

### Slide 7
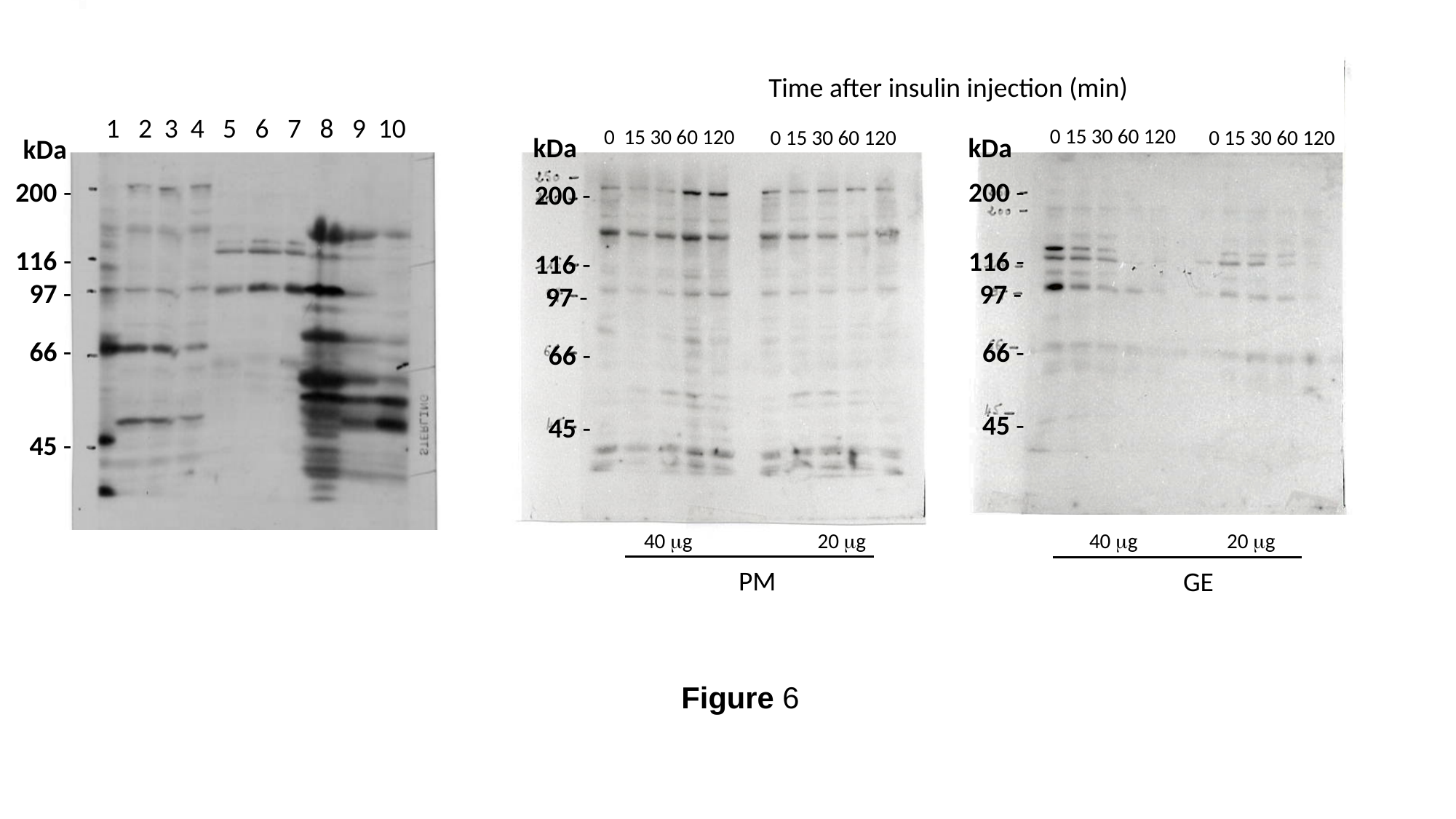

Time after insulin injection (min)
1 2 3 4 5 6 7 8 9 10
0 15 30 60 120
0 15 30 60 120
0 15 30 60 120
0 15 30 60 120
kDa
kDa
kDa
200 -
116 -
97 -
66 -
45 -
200 -
116 -
97 -
66 -
45 -
200 -
116 -
97 -
66 -
45 -
40 μg
20 μg
40 μg
20 μg
PM
GE
Figure 6

### Slide 8
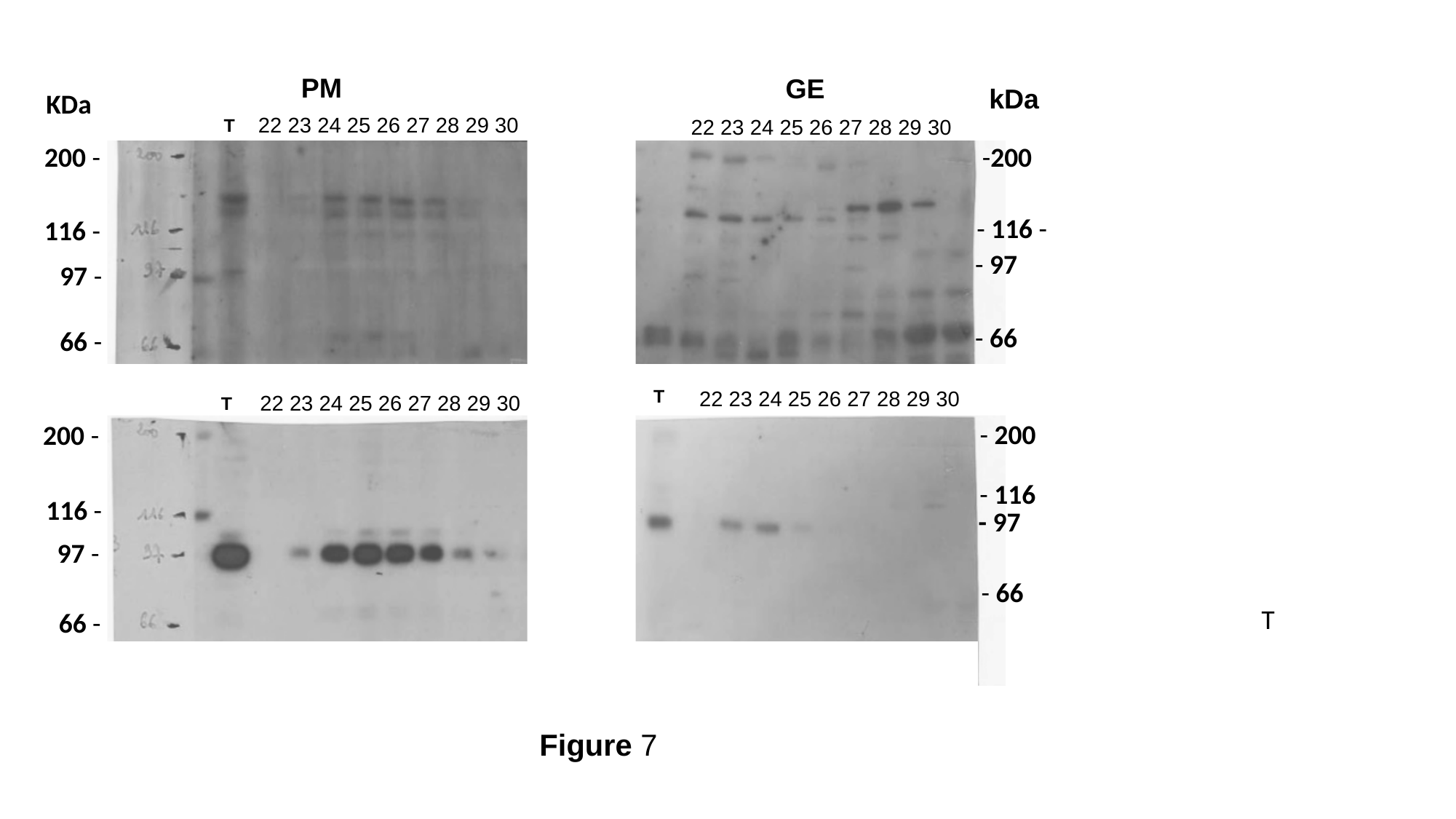

PM
GE
kDa
KDa
22 23 24 25 26 27 28 29 30
22 23 24 25 26 27 28 29 30
T
200 -
-200
- 116 -
116 -
- 97
97 -
- 66
66 -
T
22 23 24 25 26 27 28 29 30
22 23 24 25 26 27 28 29 30
T
- 200
200 -
- 116
116 -
- 97
97 -
- 66
T
66 -
Figure 7

### Slide 9
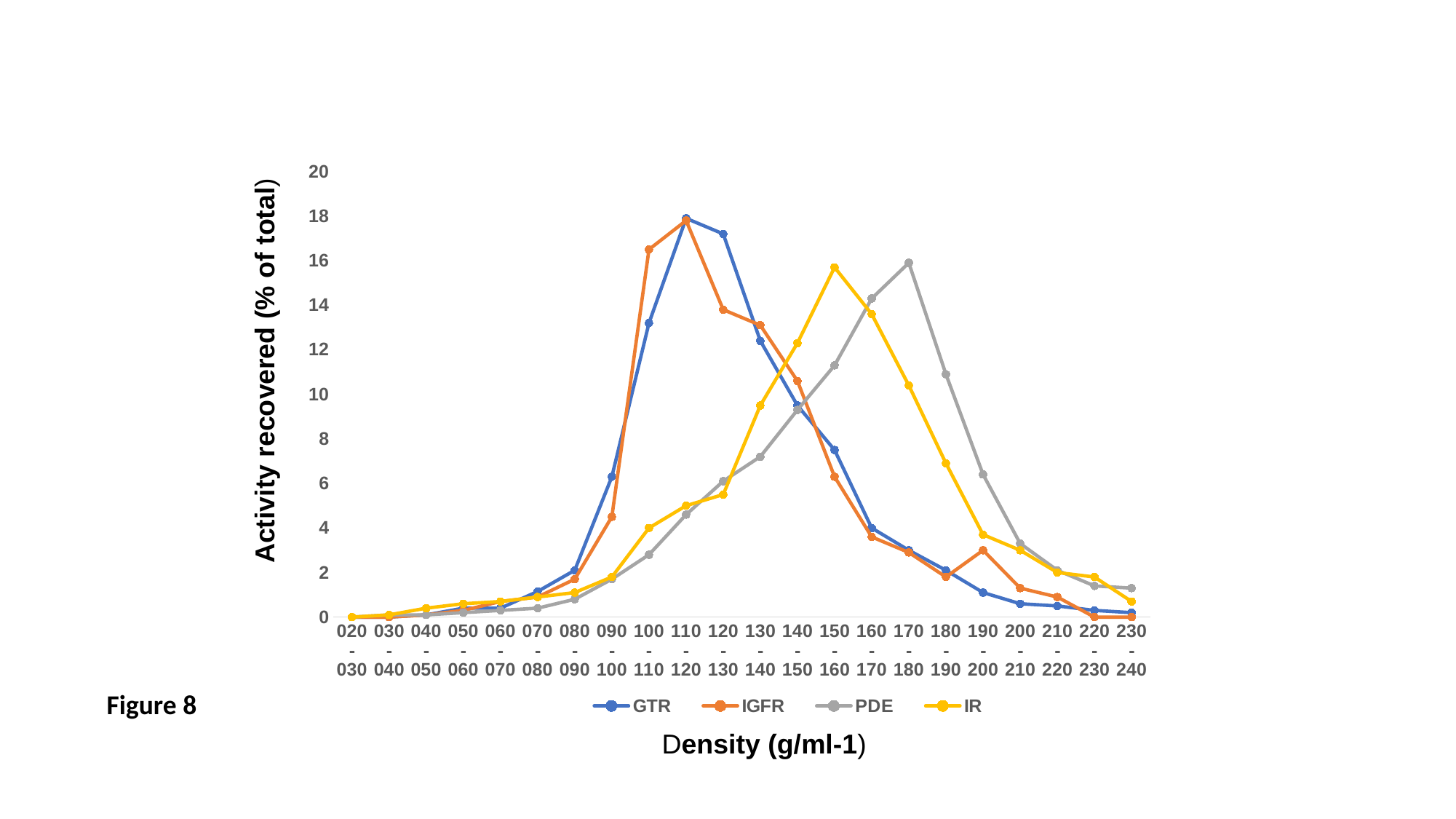

#### Chart
| Category | GTR | IGFR | PDE | IR |
|---|---|---|---|---|
| 020-030 | 0.0 | 0.0 | 0.0 | 0.0 |
| 030-040 | 0.0 | 0.0 | 0.1 | 0.1 |
| 040-050 | 0.1 | 0.1 | 0.1 | 0.4 |
| 050-060 | 0.4 | 0.3 | 0.2 | 0.6 |
| 060- 070 | 0.4 | 0.7 | 0.3 | 0.7 |
| 070-080 | 1.15 | 0.9 | 0.4 | 0.9 |
| 080-090 | 2.1 | 1.7 | 0.8 | 1.1 |
| 090-100 | 6.3 | 4.5 | 1.7 | 1.8 |
| 100-110 | 13.2 | 16.5 | 2.8 | 4.0 |
| 110-120 | 17.9 | 17.8 | 4.6 | 5.0 |
| 120-130 | 17.2 | 13.8 | 6.1 | 5.5 |
| 130-140 | 12.4 | 13.1 | 7.2 | 9.5 |
| 140-150 | 9.5 | 10.6 | 9.3 | 12.3 |
| 150-160 | 7.5 | 6.3 | 11.3 | 15.7 |
| 160-170 | 4.0 | 3.6 | 14.3 | 13.6 |
| 170-180 | 3.0 | 2.9 | 15.9 | 10.4 |
| 180-190 | 2.1 | 1.8 | 10.9 | 6.9 |
| 190-200 | 1.1 | 3.0 | 6.4 | 3.7 |
| 200-210 | 0.6 | 1.3 | 3.3 | 3.0 |
| 210-220 | 0.5 | 0.9 | 2.1 | 2.0 |
| 220-230 | 0.3 | 0.0 | 1.4 | 1.8 |
| 230-240 | 0.2 | 0.0 | 1.3 | 0.7 |Activity recovered (% of total)
Figure 8
Density (g/ml-1)
